# Macrophages sense ECM mechanics and growth factor availability through cytoskeletal remodeling to regulate their tissue repair program

**DOI:** 10.1101/2023.06.28.545586

**Authors:** Matthew L. Meizlish, Yoshitaka Kimura, Scott D. Pope, Rita Matta, Catherine Kim, Naomi Philip, Linde Meyaard, Anjelica Gonzalez, Ruslan Medzhitov

## Abstract

Tissue resident macrophages play important roles in tissue homeostasis and repair. However, how macrophages monitor and maintain tissue integrity is not well understood. The extracellular matrix (ECM) is a key structural and organizational component of all tissues. Here, we find that macrophages sense the mechanical properties of the ECM in order to regulate a specific tissue repair program. We show that macrophage mechanosensing is mediated by cytoskeletal remodeling and can be performed in three-dimensional environments through a non-canonical, integrin-independent mechanism analogous to amoeboid migration. We find that these cytoskeletal dynamics also integrate biochemical signaling by CSF1 and ultimately regulate chromatin accessibility to control the mechanosensitive gene expression program. This study suggests a distinct mode of ECM mechanosensing and growth factor signaling through which macrophages may regulate tissue repair and fibrosis.

## Introduction

The extracellular matrix (ECM) is an organized assembly of proteins and polysaccharides that is produced by cells and in turn forms the physical environment in which cells reside. The ECM is a fundamental feature of multicellular organisms, providing structural support and organization for cells within the tissue and conferring the particular mechanical properties that are necessary for the proper function of that tissue.^1–3^ The lungs, for instance, must be compliant and elastic to breathe, while bone must be able to resist large forces in order to bear the body’s weight and maintain its structure. The composition of the ECM—the repertoire of ECM components, their concentrations, and their spatial arrangement—gives rise to the particular mechanical features of each tissue.^4^

The composition and mechanical properties of the ECM can also undergo dramatic changes within a given organ. During tissue repair, activated fibroblasts called myofibroblasts deposit ECM components to reconstruct the injured tissue. This reparative phase must be followed by a phase of resolution, in which ECM deposition is suppressed and the ECM is remodeled, in order to return the tissue to a functional state. If repair instead persists and ECM deposition continues, it can lead to the development of fibrosis—dense, stiff scar tissue that replaces normal, functioning tissue, ultimately leading to organ failure.^5, 6^ Given its critical role in tissue organization, the ECM must be actively monitored, in order to maintain it in the appropriate state under normal conditions and in order to regulate the progression of tissue repair.

Macrophages play an important role in maintaining tissue homeostasis by sensing and regulating a number of homeostatic variables, like oxygen tension and osmolarity.^7–9^ Macrophages also play a key role in tissue repair. After tissue injury, monocytes produced in the bone marrow enter the tissue from the blood,^10, 11^ where they encounter the growth factor Colony-stimulating factor 1 (CSF1), which is produced by stromal cells like fibroblasts and which induces monocyte differentiation into macrophages.^12–15^ These macrophages drive ECM deposition by fibroblasts during repair^10, 16–21^ and can directly degrade and remodel ECM during resolution.^22–27^ However, it remains unknown how these cellular activities are controlled. We hypothesized that macrophages sense the state of the ECM by monitoring its chemical and/or mechanical properties.

There are several reasons that macrophages may have evolved to sense ECM mechanics: 1) The mechanical characteristics of the ECM are an emergent property of the many individual ECM components, so measuring ECM mechanics is an efficient way to assess the state of the ECM as a whole; 2) the mechanical properties of the ECM are essential to its proper function; and 3) ECM stiffness changes dramatically during tissue repair and may serve as a proxy to detect the progression of repair and avoid fibrosis.

A number of cell types are well established as mechanosensors of the ECM, including mesenchymal stem cells, fibroblasts, epithelial cells, endothelial cells, and smooth muscle cells.^28–32^ The best known model of ECM mechanosensing, established through studies of these cell types, involves adhesion to the ECM via a family of cell-surface receptors known as integrins.^33–36^ Integrin receptors are coupled intracellularly to the actin cytoskeleton. In order for the cell to sense the stiffness of the ECM, the motor protein non-muscle myosin II must pull on the actin cytoskeleton, which in turn exerts tension on integrin receptors, which pull against their ECM ligands.^28, 29, 33, 35, 37, 38^

This integrin-dependent model of ECM mechanosensing has been established through studies of cell types, like fibroblasts, that are typically stationary within tissues and that form firm adhesions to the ECM. When they do migrate, these cells must pull themselves along by exerting tension on their surroundings (‘crawling’), using integrin-based adhesions to the ECM and non-muscle myosin II—the same mechanisms that they use for mechanosensing. This mode of migration is known as mesenchymal migration.

In contrast, while leukocytes can utilize integrin-based adhesions and do require integrins to adhere to two-dimensional (2D) surfaces, they do not require integrin receptors in order to migrate within three-dimensional (3D) tissues.^39^ Instead of attaching to the ECM and crawling, leukocytes are able to migrate in an adhesion-independent fashion, driven instead by actin protrusion at the leading edge of the cell, a mode of migration that resembles ‘swimming’. This allows them to infiltrate a diverse range of tissues, migrate outside of prescribed paths, and achieve much higher speeds than is possible through the traditional, crawling mode of migration.^39–41^ This mode of migration is known as ameboid migration.

Recent studies have revealed that macrophages on 2D surfaces can utilize the canonical mode of adhesion-dependent mechanosensing to detect substrate stiffness and activate established downstream mechanisms like the mechanosensitive transcription factor YAP.^38, 42^ Mechanosensitive ion channels, which play a key role in responding to transient mechanical stimuli like touch in sensory neurons,^43, 44^ have also been implicated in macrophage mechanosensing. PIEZO1 mediates macrophage responses to cyclic pressure in the lungs, and there is evidence that PIEZO1 and TRPV4 are involved in sensing substrate stiffness on 2D surfaces, where they likely complement the adhesive function of integrins.^45–49^ However, little is known about how macrophages in 3D tissues sense the properties of the ECM.

Here, we find that macrophages in 3D environments can sense the mechanical properties of the ECM through a cytoskeleton-dependent but integrin-independent mechanism that resembles their mode of migration and is distinct from the canonical model of mechanosensing elucidated in strictly adhesive cell types like fibroblasts. This macrophage mechanosensing pathway controls a specific gene expression program involved in tissue repair. Interestingly, signaling by the growth factor CSF1 also remodels the cytoskeleton to converge on the same gene expression program.

We find that this cytoskeletal remodeling ultimately alters chromatin accessibility in a site-specific fashion to regulate gene expression. Finally, we find *in vivo* that macrophages exhibit these same mechanosensitive responses at the cytoskeletal and transcriptional levels as tissue stiffness increases over the course of tissue repair. Altogether, this study identifies a specific tissue repair program in macrophages that is regulated by an integrin-independent mechanosensing mechanism.

## Results

### Macrophages sense ECM mechanics to control tissue repair-associated gene expression

In order to model a simple, well-defined tissue environment, we cultured bone marrow-derived macrophages (BMDMs) within three-dimensional (3D) hydrogels composed of type I collagen, the most abundant ECM protein in most tissues (**Fig. 1A**). To determine whether macrophages can sense changes in the ECM, we varied the concentration of collagen from 2 mg/mL (low collagen) to approximately 7 mg/mL (high collagen), which also altered the stiffness of the collagen gels (**Fig. S1A)** and the pore size of the fibrillar network (**Fig. S1B-C**).

**Fig. 1.**
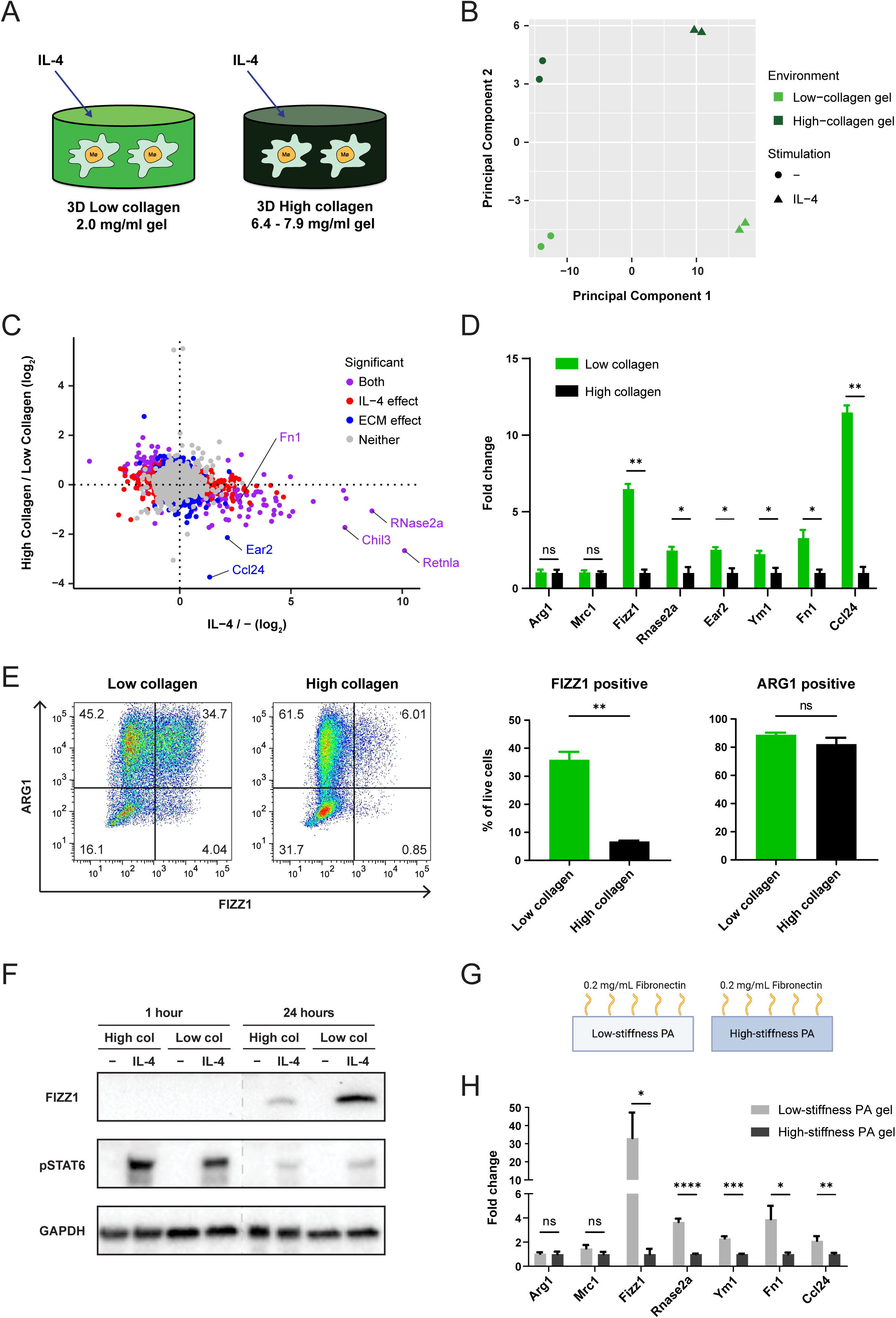
Sensing of ECM stiffness controls a tissue repair-associated gene expression program in macrophages. **(A)** Schematic overview of the experimental setup for macrophage culture within 3D collagen hydrogels. **(B and C)** Principal component analysis (B) and plot of all individual genes (C) in RNA-seq data from BMDMs cultured in low- or high-collagen gels with or without IL-4. Plots in (C) are colored according to the statistical significance of gene expression changes caused by culture in high-vs. low-collagen gels, as well as treatment with IL-4 vs. control (q-value < 0.2). Retnla = Fizz1, Chil3 = Ym1. **(D)** Fold-change of relative expression of selected genes in BMDMs cultured in low-collagen gels compared to high-collagen gels. Mean gene expression in high-collagen gels is shown as 1. **(E)** FIZZ1 and ARG1 protein expression in BMDMs cultured in low- or high-collagen gels with IL-4, analyzed by flow cytometry following the gating strategy displayed in Fig. S1E. Representative plots (left) and the percentage of FIZZ1-positive and ARG1-positive cells (right) are shown. **(F)** Western blot showing FIZZ1 and phospho-STAT6 (pSTAT6) protein 1 hour and 24 hours after control or IL-4 treatment in low- or high-collagen gels. **(G)** Schematic overview of the experimental setup for polyacrylamide-fibronectin (PA) hydrogel culture. **(H)** Fold-change of relative gene expression in BMDMs cultured on low-stiffness PA compared to high-stiffness PA gels. Mean gene expression on high-stiffness gels is shown as 1. Data are represented as mean ± SD. Multiple t test (D and H) and Student’s t test (E) were used for statistical analysis. *p < 0.05, ** p < 0.01, *** p < 0.001, **** p < 0.0001. ns; not significant.

IL-4 induces a well-defined transcriptional program in macrophages to orchestrate a tissue-reparative response.^50^ We hypothesized that the transcriptional response to IL-4 may depend on the state of the extracellular matrix, in order to appropriately regulate repair. Indeed, we found that macrophages cultured in high-collagen compared to low-collagen gels showed global differences in IL-4 induced gene expression (**Fig. 1B**), suggesting that macrophages can sense and respond to the state of the extracellular matrix and integrate this information with the instructive signals provided by cytokines.

Turning to a gene-based analysis, we identified a cluster of genes that were induced by IL-4 and suppressed in high-collagen compared to low-collagen gels (**Fig. 1C**). Several of these genes, including *Fizz1* (*Retnla*, RELMα), *Rnase2a*, *Ear2*, *Ym1* (*Chil3*), *Fn1*, and *Ccl24*, were consistently expressed at higher levels in low-collagen gels compared to high-collagen gels, while other genes induced by IL-4, such as *Arg1* and *Mrc1*, were consistently unaffected by the state of the ECM (**Fig. 1D**). *Fizz1*, one of the most highly regulated genes, is known to play a critical role in tissue repair in the lung and in the skin.^21, 51^ It is secreted by macrophages and is thought to act on fibroblasts to regulate myofibroblast differentiation, collagen production, and collagen cross-linking.^21, 52^

Expression of these ECM-sensitive genes, represented by *Fizz1*, showed a dose response to the concentration of collagen, while ECM-insensitive genes, represented by *Arg1*, did not (**Fig. S1D**). We confirmed these findings on the protein level by performing flow cytometry with intracellular protein staining (**Fig. S1E**). We found that FIZZ1 protein levels were markedly suppressed in high-collagen compared to low-collagen gels, while ARG1 protein levels were unaffected by the state of the ECM (**Fig. 1E**).

The identification of two distinct patterns of IL-4-induced gene expression suggested that the effects of the ECM were not mediated by changing the strength of IL-4 signaling. Indeed, phosphorylation of STAT6, the major mediator of IL-4 signaling, was equivalent or even slightly increased in high-compared to low-collagen gels, while FIZZ1 protein expression was markedly reduced in high-collagen gels (**Fig. 1F**). These data indicate that there are distinct sub-programs induced by IL-4, which can be regulated independently. At least one of those programs, represented by *Fizz1*, is sensitive to changes in the tissue environment and in particular to the extracellular matrix.

Next, we wanted to determine which specific features of the ECM are detected by macrophages. In collagen gels, the concentration of collagen is directly related to the mechanical properties of the hydrogel (**Fig. S1A-C**). While this accurately reflects the relationship that exists *in vivo* as ECM accumulates, it limits our ability to experimentally dissect which properties of the ECM are monitored. In order to decouple ECM concentration from ECM mechanics, we cultured macrophages on polyacrylamide (PA) hydrogels conjugated to the major ECM protein fibronectin (**Fig. 1G**). In this system, the stiffness of the hydrogel can be controlled by varying the density of polyacrylamide crosslinking, while the PA network is inert and not bound by cells. The cells bind to the conjugated fibronectin, the concentration of which is held constant.^28, 53^ Thus, the effect of altering ECM stiffness can be isolated. When we cultured macrophages on high-stiffness compared to low-stiffness PA-Fibronectin gels (**Fig. S1F**) the ECM-sensitive gene expression program was suppressed (**Fig. 1H**), similar to what we observed in high-collagen compared to low-collagen gels. Meanwhile, ECM-independent genes like Arg1 were again unaffected (**Fig. 1H**). These data establish that macrophages can sense ECM stiffness and that the ECM-sensitive program that we have identified is in fact a mechanosensitive gene expression program.

### Macrophage mechanosensing is mediated by cytoskeletal remodeling

To dissect the mechanism of macrophage mechanosensing within tissues, we utilized the 3D collagen gel system. Using live cell imaging, we observed that macrophages in low-collagen gels migrate more rapidly and over a greater distance than macrophages in high-collagen gels (**Movie. S1**, **Fig. 2A**). They also had marked differences in morphology: In low-collagen gels, macrophages were typically round or had short protrusions that rapidly turned over, while in high-collagen gels they were often dendritic, with numerous long actin protrusions that retracted more slowly (**Movie. S1**). We further visualized these morphologic differences by phalloidin staining of actin filaments, followed by confocal microscopy (**Fig. 2B**) and quantified them by cell sphericity (**Fig. 2C**). We hypothesized that these cytoskeletal dynamics may serve as an intracellular measure of the extracellular mechanical environment and may ultimately control the observed changes in gene expression.

**Fig. 2.**
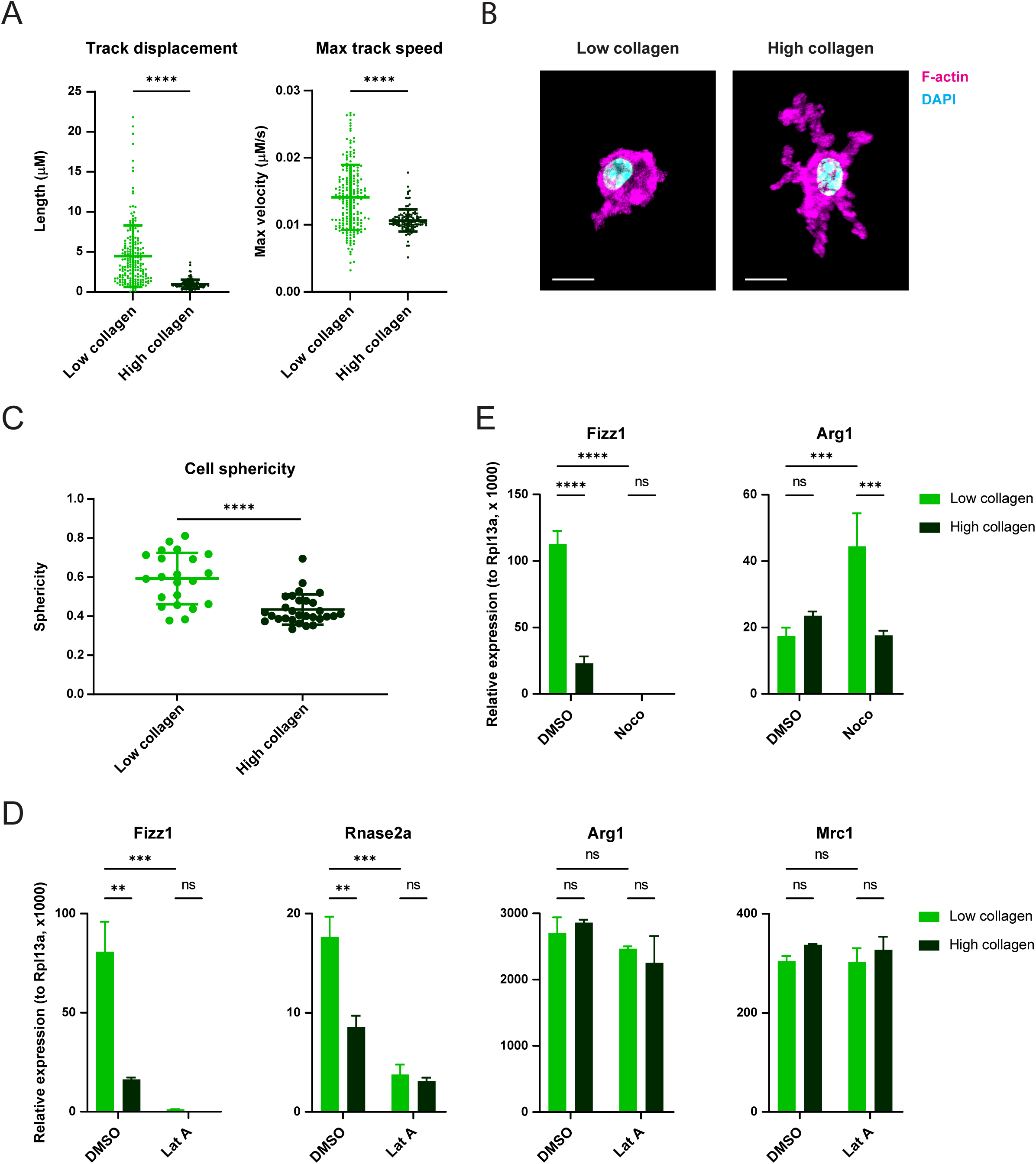
Cytoskeletal remodeling is required for macrophage mechanosensing. **(A)** Quantification of migration distance and speed of BMDMs cultured in low- or high-collagen gels after stimulation with IL-4. **(B and C)** Representative confocal images (B) and quantification of the sphericity (C) of BMDMs cultured in low- or high-collagen gels and stimulated with IL-4. Cells were stained with phalloidin (magenta) to visualize F-actin and DAPI (blue) to visualize the nucleus. Scale bar = 10 μm. **(D)** Relative gene expression in IL-4-stimulated BMDMs cultured in low- or high-collagen gels with DMSO or latrunculin A (Lat A). **(E)** Relative gene expression in IL-4-stimulated BMDMs cultured in low- or high-collagen gels with DMSO or nocodazole (Noco). Data are represented as mean ± SD. Student’s t test (A and C) and two-way ANOVA (D and E) are used for statistical analysis. ** p < 0.01, *** p < 0.001, **** p < 0.0001. ns; not significant.

To test this hypothesis, we used chemical inhibitors to directly manipulate the macrophage cytoskeleton. When macrophages were treated with latrunculin A (LatA) to inhibit actin polymerization, mechanosensitive gene expression was potently suppressed, while non-mechanosensitive genes like *Arg1* and *Mrc1* were unaffected (**Fig. 2D**). Similarly, disrupting microtubule dynamics with nocodazole profoundly suppressed mechanosensitive genes like *Fizz1*, unlike other IL-4-induced genes like *Arg1* (**Fig. 2E**). Furthermore, in the presence of latrunculin A (**Fig. 2D**) or nocodazole (**Fig. 2E**), mechanosensitive genes were no longer responsive to changes in the ECM (low collagen vs. high collagen), indicating that intact cytoskeletal dynamics are required for macrophage mechanosensing. Altogether, these data show that macrophage mechanosensing is mediated by cytoskeletal responses to the tissue environment and that these cytoskeletal dynamics specifically regulate the mechanosensitive sub-program of the macrophage IL-4 response.

### Macrophages perform integrin-independent mechanosensing

We next investigated whether integrin-based interactions with the ECM are required for macrophage mechanosensing, as would be expected for mesenchymal cell types like fibroblasts. We first asked whether fibroblasts follow the expected biological rules in this experimental system. We found that murine embryonic fibroblasts (MEFs) upregulated established mechanosensitive genes *Ctgf* and αSMA (*Acta2*) in high-compared to low-collagen gels (**Fig. S2A**). We confirmed that fibroblast mechanosensing in this setting requires non-muscle myosin II (using the inhibitor blebbistatin, **Fig. S2A**) and β1 integrin (using a β1 integrin blocking antibody, **Fig. S2B**). These results are consistent with the prevailing model that fibroblast mechanosensing of the ECM requires integrin-based adhesion and mechanical tension exerted by non-muscle myosin II.

We then investigated whether macrophage mechanosensing is accomplished by the same mechanisms as adhesive cell types or in an integrin-independent fashion, analogous to leukocyte migration in 3D tissues.^39–41^ Interestingly, both bone marrow-derived macrophages and tissue-resident macrophages show minimal expression of canonical collagen-binding integrins, which are heterodimers composed of the β1 integrin chain paired with the α1, α2, α10, or α11 integrin chain (**Fig. S3**).^54–56^ When we inhibited the function of all of these collagen-binding pairs using the β1 integrin blocking antibody, we found that it had no effect on mechanosensitive gene expression (**Fig. 3A**). Blocking β2 integrin—the β chain for all of the ‘leukocyte integrins’, which are classically involved in cell-cell interactions but some of which have also been reported to interact with collagen—also had no effect on ECM sensing (**Fig. 3B**).^57, 58^ To further test the involvement of integrins, we performed siRNA knockdown of *Talin1*, an adapter protein that connects integrins to the cytoskeleton and is required for nearly all integrin activity. Though *Talin1* expression was successfully suppressed, there was no effect on *Fizz1* expression or responsiveness to the ECM (**Fig. 3C**), further indicating that macrophage mechanosensing in 3D collagen gels is integrin-independent. Interestingly, inhibiting non-muscle myosin II with blebbistatin suppressed *Fizz1* expression, but it remained sensitive to changes in the ECM even after blebbistatin treatment (**Fig. 3D**). These data indicate that macrophage mechanosensing in 3D collagen gels is cytoskeleton-dependent but both integrin- and myosin-II-independent, unlike canonical mechanosensing mechanisms in fibroblasts and other adhesive cell types.

**Fig. 3.**
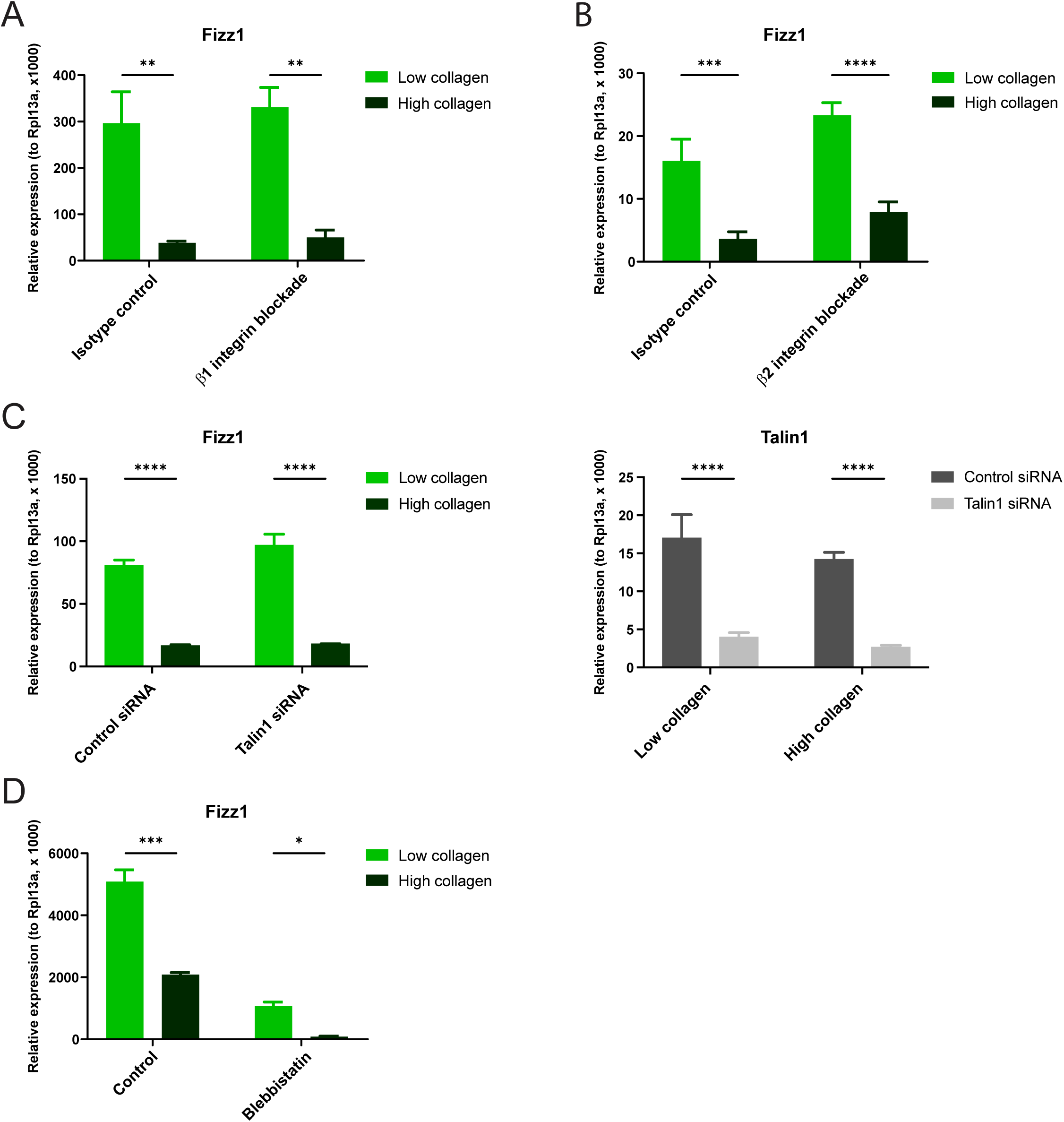
Macrophage mechanosensing does not require integrins or myosin II. **(A and B)** Relative gene expression in BMDMs treated with β1 integrin blocking antibody (A), β2 integrin blocking antibody (B), or isotype-matched controls (A and B) and cultured in low- or high-collagen gels in the presence of IL-4. **(C)** Relative gene expression in BMDMs treated with control siRNA or Talin1 siRNA and cultured in low- or high-collagen gels in the presence of IL-4. **(D)** Relative gene expression in BMDMs treated with control or blebbistatin and cultured in low- or high-collagen gels in the presence of IL-4. Data are represented as mean ± SD. Two-way ANOVA was used for statistical analysis. *p < 0.05, ** p < 0.01, *** p < 0.001, **** p < 0.0001.

We also tested other mechanisms that have been implicated in macrophage mechanosensing on 2D surfaces.^42, 45, 46, 49^ We did not detect appreciable YAP expression in BMDMs (**Fig. S4A**) and found that inhibiting the downstream transcription factor TEAD did not affect macrophage mechanosensing in 3D collagen gels (**Fig. S4B**). Investigating mechanosensitive ion channels, we found that deleting PIEZO1 did not change *Fizz1* expression (**Fig. S4C**), and neither inhibiting TRPV4 nor the class of mechanosensitive ion channels blunted macrophage responses to ECM (**Fig. S4D**). While these mechanisms may play an important role in integrin-dependent mechanosensing on 2D surfaces, they do not appear to mediate integrin-independent mechanosensing by macrophages within 3D tissues.

### Macrophage growth factor CSF1 remodels the cytoskeleton to regulate the mechanosensitive gene expression program

Macrophages sense many other features of the tissue environment, in addition to ECM mechanics.^7, 9^ Among the most important regulators of macrophage biology is the lineage-restricted growth factor CSF1, which stimulates monocyte-to-macrophage differentiation, as well as macrophage survival and proliferation. CSF1 is produced by other cell types within the tissue, such as fibroblasts, and it can serve as an indicator of fibroblast cell number, which changes dynamically during tissue repair.^12–15^ To our surprise, we found that stimulation of macrophages with recombinant CSF1 specifically suppressed the mechanosensitive gene expression program that we had identified, similar to high-stiffness environments, while it increased or had no effect on the expression of non-mechanosensitive IL-4-induced genes like Arg1 and Mrc1 (**Fig. 4A**).

**Fig. 4.**
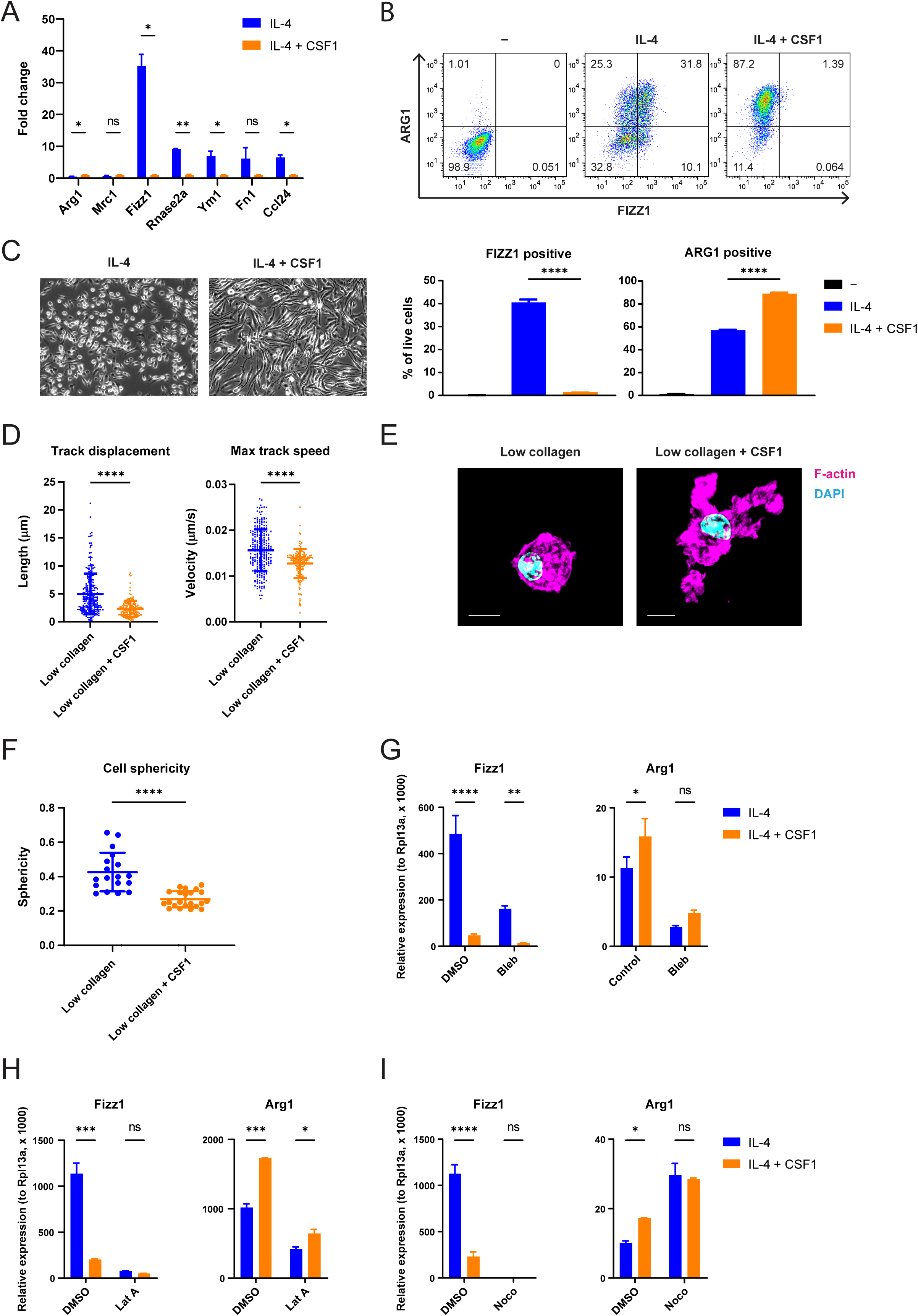
CSF1 sensing by macrophages induces cytoskeletal remodeling to regulate the mechanosensitive gene expression program. **(A)** Fold-change of relative gene expression in BMDMs cultured on tissue culture (TC) plates and stimulated with IL-4 only, compared to both IL-4 and CSF1. Mean gene expression in cells treated with both IL-4 and CSF1 is shown as 1. **(B)** FIZZ1 and ARG1 protein expression in BMDMs cultured on TC plates and left unstimulated, stimulated with IL-4 only, or stimulated with both IL-4 and CSF1. Protein expression was analyzed by flow cytometry following the gating strategy displayed in Fig. S1E. Representative plots (upper) and the percentage of FIZZ1-positive and ARG1-positive cells (lower) are shown. **(C)** Representative microscopic images of BMDMs cultured on TC plates in the presence of IL-4 with or without CSF1. **(D)** Quantification of migration distance and speed of BMDMs cultured in low-collagen gels in the presence of IL-4 with or without CSF1. **(E and F)** Representative confocal images (E) and quantification of the sphericity (F) of BMDMs cultured in low-collagen gels in the presence of IL-4 with or without CSF1. Cells were stained with phalloidin (magenta) to visualize F-actin and DAPI (blue) to visualize the nucleus. Scale bar = 10 μm. **(G)** Relative gene expression in IL-4-stimulated BMDMs cultured on TC plates and treated with control or CSF1 in the presence or absence of blebbistatin. **(H)** Relative gene expression in IL-4-stimulated BMDMs cultured on TC plates and treated with control or CSF1 in the presence or absence of latrunculin A (Lat A). **(I)** Relative gene expression in IL-4-stimulated BMDMs cultured on TC plates and treated with control or CSF1 in the presence or absence of nocodazole (Noco). Data are represented as mean ± SD. Student’s t test (A, D, and F), one-way ANOVA (B), two-way ANOVA (G, H and I) are used for statistical analysis. *p < 0.05, ** p < 0.01, *** p < 0.001, **** p < 0.0001. ns; not significant.

We confirmed these findings for FIZZ1 and ARG1 on the protein level by flow cytometry (**Fig. 4B**). We also observed that CSF1 induces an elongated macrophage morphology on tissue culture plates (**Fig. 4C**). We then performed live cell microscopy and confocal imaging in 3D collagen gels, which showed that macrophages treated with CSF1 become highly dendritic in 3D environments, migrating more slowly and extending long actin protrusions (**Movie. S2**, **Fig. 4D-F**), similar to the macrophage behavior we observed in high-collagen gels. Thus, CSF1 and high-stiffness ECM have similar effects on both macrophage cytoskeletal dynamics and gene expression.

Because we found that the effects of ECM mechanics on macrophage gene expression are cytoskeleton-dependent, we reasoned that CSF1 may also act through its effects on the cytoskeleton in order to control the mechanosensitive gene expression program. To test this hypothesis, we treated macrophages with CSF1 under normal conditions or after pharmacologically disrupting the cytoskeleton. Consistent with our findings for macrophage mechanosensing, the effect of CSF1 on *Fizz1* was not dependent on non-muscle myosin II (**Fig. 4G**). However, the effect of CSF1 on *Fizz1* expression was dependent on both actin (**Fig. 4H**) and microtubule dynamics (**Fig. 4I**). In contrast, the induction of Arg1 by CSF1 was actin-independent (**Fig. 4H**), though interestingly was abrogated in the presence of nocodazole (**Fig. 4I**). Altogether, we find that CSF1 signaling, like ECM mechanosensing, acts through cytoskeletal remodeling to regulate a specific tissue repair program in macrophages that is sensitive to changes in the tissue environment.

### Cytoskeletal remodeling controls site-specific chromatin accessibility

Next, we investigated how cytoskeletal remodeling in macrophages ultimately controls transcriptional regulation. To determine whether changes in actin dynamics result in changes in chromatin accessibility, we performed ATACseq in BMDMs treated with or without IL-4 and with or without latrunculin A and CSF1, both of which suppress expression of *Fizz1* by acting on the macrophage cytoskeleton. We found that treatment with IL-4 induces chromatin accessibility at the critical region for IL-4-induced activation within the *Fizz1* promoter and that this chromatin opening is potently suppressed by both cytoskeletal perturbations (**Fig. 5A**).^59^ In contrast, the *Arg1* locus did not show notable changes in chromatin accessibility, including at the enhancer required for IL-4-induced expression (**Fig. 5B**).^60, 61^ These data indicate that macrophage cytoskeletal dynamics regulate chromatin accessibility in a site-specific fashion that aligns with the changes in gene expression we observed.

**Fig. 5.**
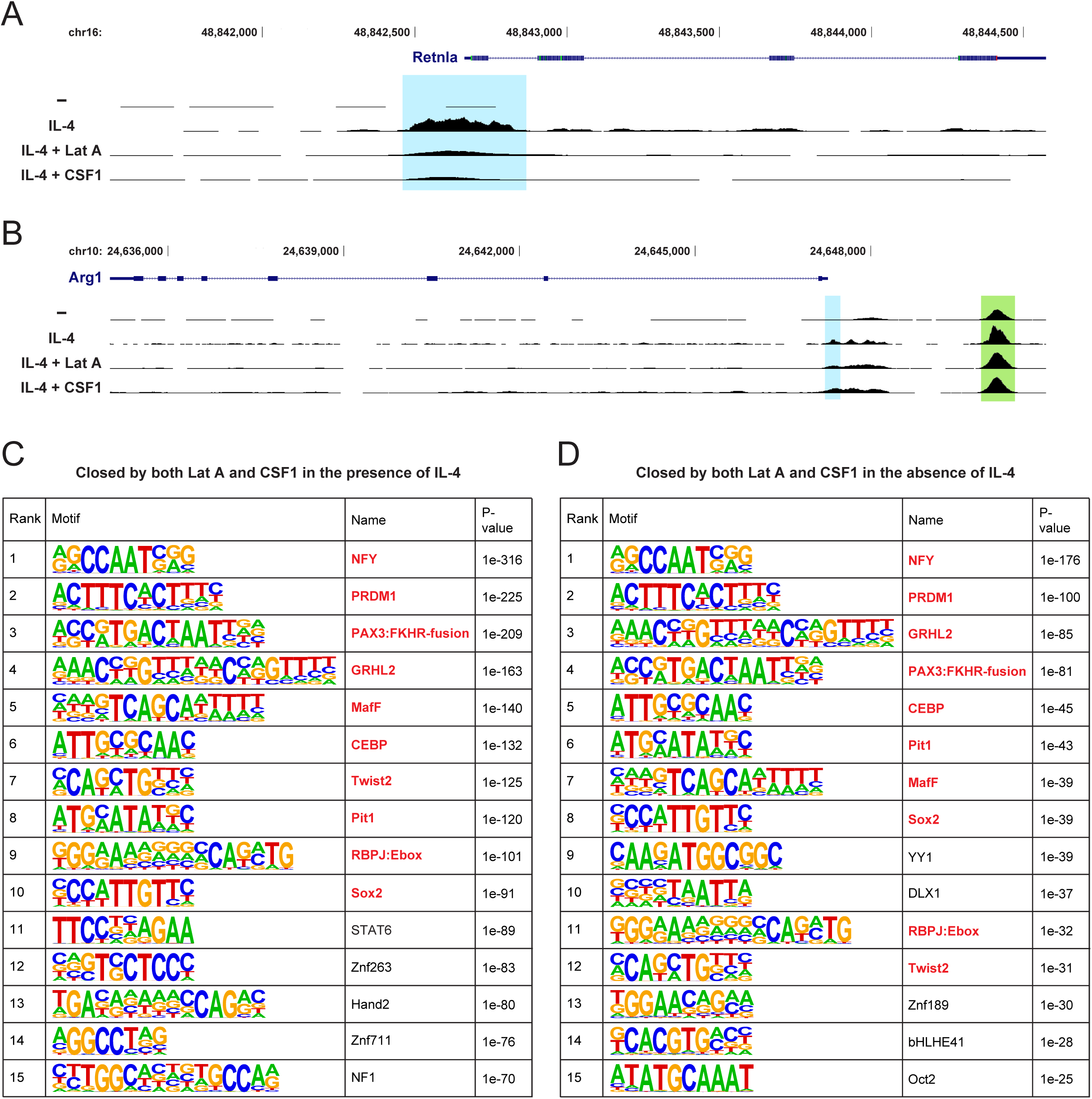
Cytoskeletal remodeling controls chromatin accessibility in a site-specific fashion. **(A-B)** ATAC-seq signal in genomic regions upstream of and including the *Retnla* (*Fizz1*, A) and *Arg1* (B) genes, analyzed in BMDMs cultured on TC plates and treated with control, IL-4, IL-4 and Lat A, or IL-4 and CSF1. *Retnla* and *Arg1* promoter regions are highlighted by blue rectangles. Known *Arg1* enhancer region is highlighted by a green rectangle. **(C-D)** Transcription factor-binding motifs enriched at genomic sites closed by both Lat A and CSF1 in the presence (C) or absence (D) of IL-4. The top 15 motifs, ranked by statistical significance, are shown. The shared motifs between (C) and (D) are highlighted as red.

We then took a global view of the changes in the chromatin landscape induced by these cytoskeletal perturbations. First, we asked which transcription factor motifs were enriched in regions of chromatin that, like *Fizz1*, were closed by LatA and CSF1 in the context of IL-4 stimulation (**Fig. 5C**). Several motifs were highly enriched, including C/EBP and STAT6, which bind to the *Fizz1* promoter to activate *Fizz1* expression.^59^ We then analyzed the chromatin regions closed by LatA and CSF1 in the absence of IL-4 stimulation (untreated; **Fig. 5D**). Interestingly, we found a highly overlapping set of enriched motifs. The top 10 transcription factor motifs (ranked by statistical significance) in the IL-4 context also appeared in the top 15 motifs in the untreated context. These results suggest that cytoskeletal remodeling may regulate particular regions of chromatin, preparing the chromatin landscape for—and modulating the response to— additional signals, such as IL-4, that trigger transcriptional activation. These data also propose a candidate set of transcription factors that may be involved in mechanosensitive gene regulation in macrophages across biological contexts.

### Macrophages exhibit cytoskeletal remodeling and mechanosensitive gene regulation as ECM accumulates during tissue repair

Finally, we tested the hypothesis that mechanosensing allows macrophages to respond to changes in tissue stiffness *in vivo* during the course of tissue repair. Using a bleomycin lung injury model, we observed a progressive increase in ECM deposition (**Fig. 6A**), and collagen in particular (**Fig. 6B**), in the days after the initial injury, leading to tissue fibrosis by day 12, as expected.^62^ Consistent with our findings in bioengineered hydrogels, this increase in tissue stiffness over time was accompanied by suppression of *Fizz1* gene expression in the lung, while *Arg1* expression was unaffected (**Fig. 6C**). Using flow cytometry with intracellular protein staining, we confirmed that macrophages in the lung suppress FIZZ1 production while maintaining steady ARG1 production as tissue repair progresses (**Fig. 6D, Fig. S5**). Finally, we asked whether macrophages exhibit the same cytoskeletal responses to changing ECM mechanics in the lungs as we observed *in vitro*.

**Fig. 6.**
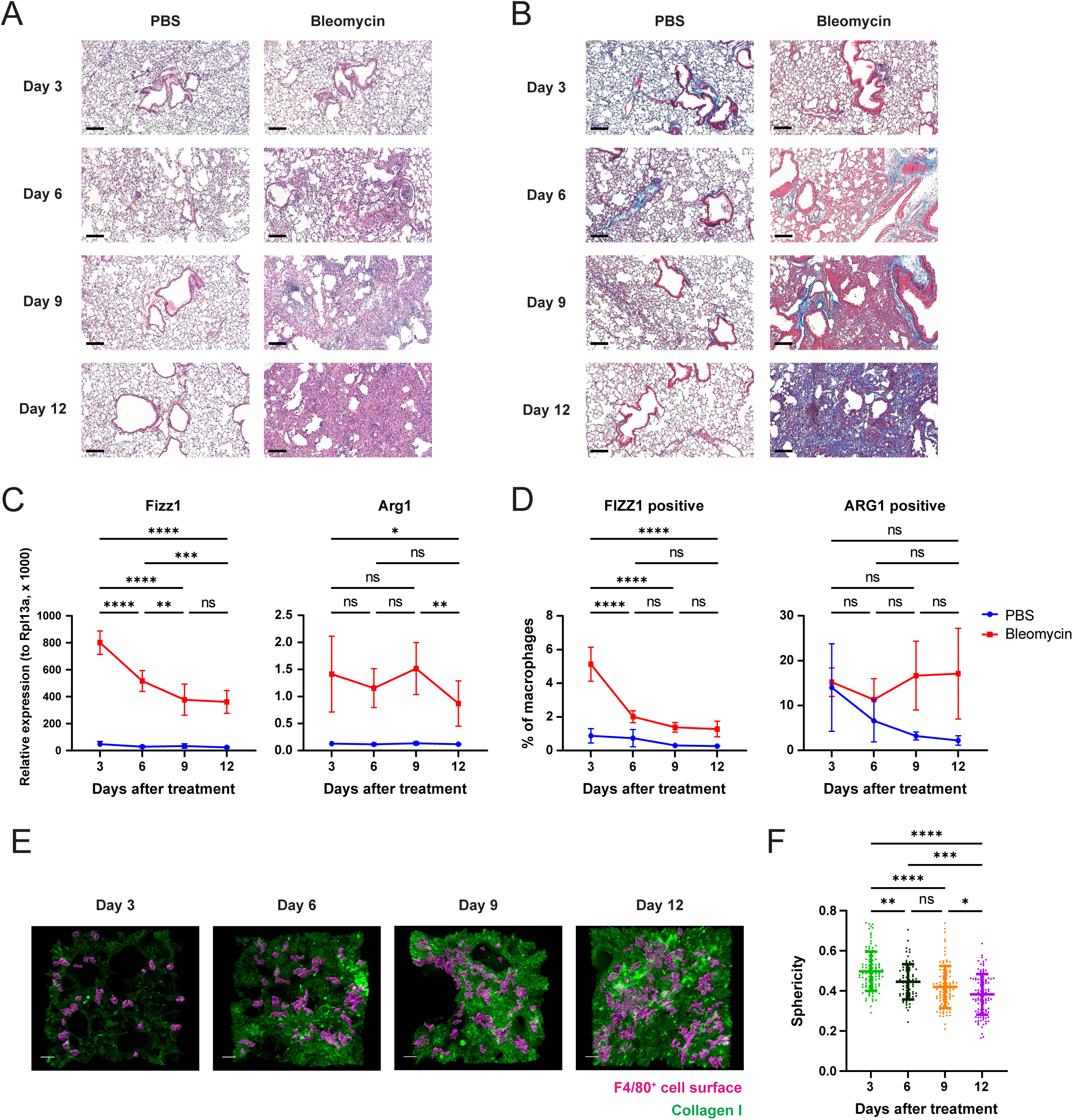
ECM deposition during tissue repair *in vivo* induces cytoskeletal remodeling and suppresses mechanosensitive gene expression in macrophages. **(A and B)** Representative microscopic images of H&E staining (A) and Masson’s trichrome staining (B) of lungs at 3, 6, 9, and 12 days after oropharyngeal PBS or bleomycin administration to mice (n = 3/group/time point). Scale bar = 100 μm. **(C)** Relative gene expression in whole lungs from PBS- or bleomycin-treated mice (n = 6-7/group/time point). Statistically significant differences between time points in the bleomycin-treated group are shown. **(D)** FIZZ1 and ARG1 protein expression in lung macrophages in PBS- or bleomycin-treated mice, analyzed by flow cytometry following the gating strategy in Fig. S5. The percentage of FIZZ1-positive and ARG1-positive cells are shown (n = 4-5/group/time point). Statistically significant differences between time points in the bleomycin-treated group are indicated. **(E)** Representative snapshots of 3D-reconstructed confocal images of lungs from bleomycin-treated mice (n = 3/time point). Surfaces of F4/80^+^ cells (magenta) and collagen I (green) are shown. Scale bar = 20 μm. **(F)** Quantification of the sphericity of F4/80+ cells in (E). Two-way ANOVA (C and D) and one-way ANOVA (F) are used for statistical analysis. *p < 0.05, ** p < 0.01, **** p < 0.0001. ns; not significant.

We stained tissue sections using antibodies against collagen I and the macrophage marker F4/80, in order to assess macrophage morphology. As collagen accumulated over time, the macrophages in the lung became less spherical and increasingly dendritic (**Fig. 6E-F**). These results are consistent with our findings in 3D collagen hydrogels, suggesting that the mechanosensing program that we identified *in vitro* is operative in complex tissues *in vivo* and is sensitive to the dynamic changes in ECM that take place during tissue repair.

## Discussion

In this study, we found that macrophages can sense the mechanical properties of their environment to regulate a specific tissue repair program. We identified an integrin-independent mechanosensing mechanism that functions through changes in intracellular cytoskeletal dynamics, which can also integrate signaling by the growth factor CSF1. We found that these cytoskeletal dynamics ultimately modify chromatin availability to regulate mechanosensitive gene expression. Finally, we saw that macrophages in the lung exhibit these mechanosensitive responses as ECM accumulates during tissue repair. Altogether, we found that the macrophage cytoskeleton integrates mechanical and biochemical information from the environment, in order to regulate a transcriptional program involved in ECM regulation and tissue repair (**Fig. 7**).

**Fig. 7.**
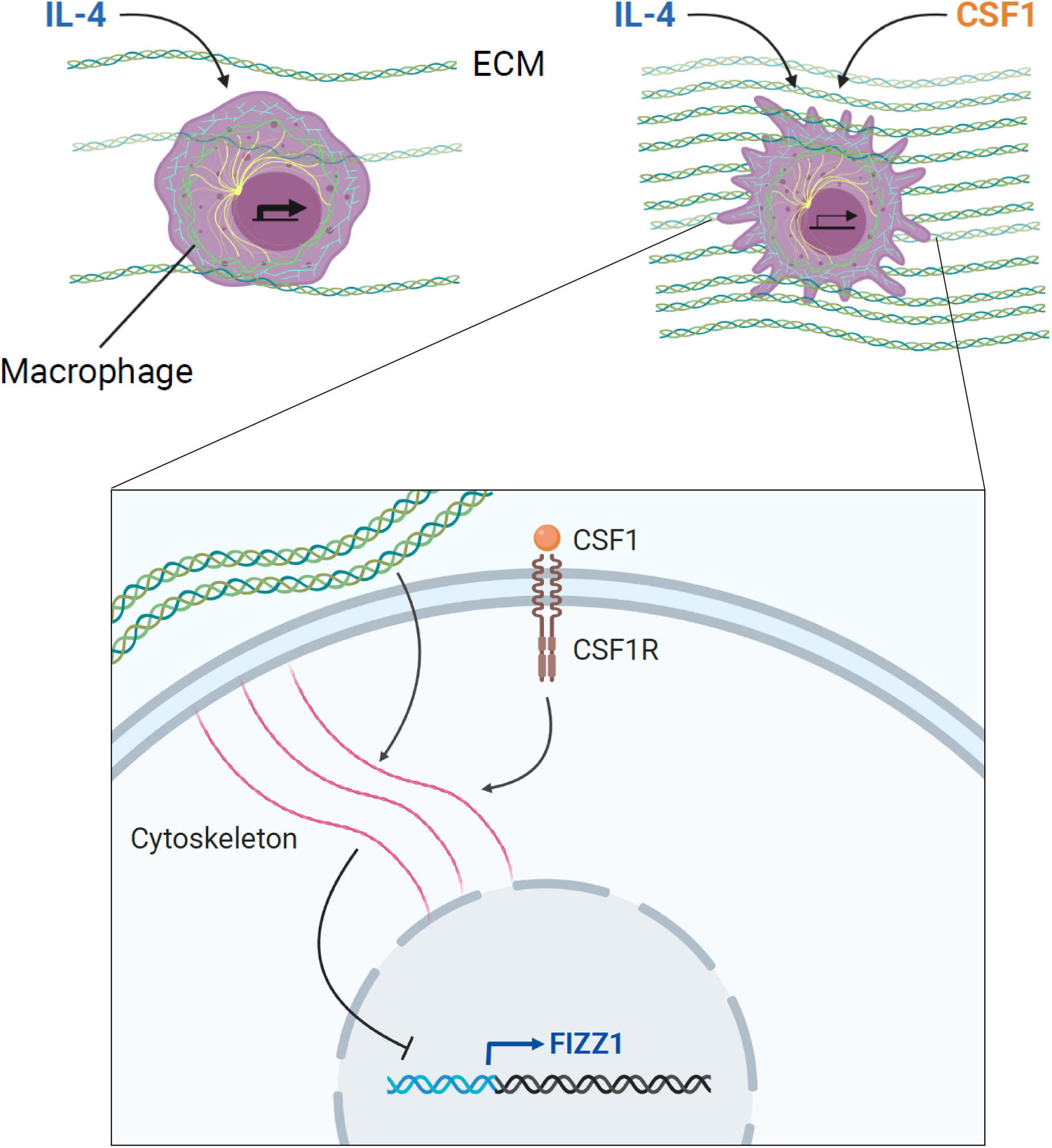
Model: ECM mechanics and CSF1 signaling are integrated through cytoskeletal remodeling to regulate tissue repair-associated gene expression in macrophages. Macrophages sense ECM mechanics through integrin-independent cytoskeletal remodeling in order to regulate the expression of a subset of tissue repair-associated genes induced by IL-4, including *Fizz1*. Signaling by CSF1, the lineage-restricted growth factor for macrophages, also converges on these cytoskeletal dynamics to regulate this mechanosensitive gene expression program. Macrophage cytoskeletal dynamics ultimately control chromatin accessibility at regulatory elements of target genes, translating environmental sensing into transcriptional regulation. These mechanisms may allow for appropriate regulation of tissue repair: As ECM stiffness and CSF1 production increase over the course of repair, macrophages undergo cytoskeletal remodeling and suppress genes that drive repair, which may be critical for resolving tissue repair and avoiding fibrosis.

Though the extracellular matrix is a fundamental element of all tissues, the logic of its regulation is poorly understood. Fibroblasts are the major producers of ECM within tissues and are well-established sensors of ECM mechanics. Paradoxically, however, when fibroblasts sense increased ECM stiffness, they respond by increasing ECM deposition and contraction, further stiffening the ECM.^63–65^ Thus, fibroblasts appear to be engaged in a positive feedback loop with respect to the ECM, which, if unrestrained, can lead to fibrosis. Yet, in most cases, ECM is maintained in the appropriate state under homeostatic conditions, and tissue repair is successfully resolved without resulting in fibrosis. These outcomes indicate that there must be a source of negative feedback on ECM deposition that has not yet been identified. We hypothesized that macrophages may act as ECM sensors to negatively regulate ECM deposition and tissue repair.

In this study, we found that macrophage sensing of increased ECM stiffness suppresses a pro-reparative gene expression program. The same program is suppressed by increased CSF1 concentration, which likely serves to report on the number of local fibroblasts, the predominant producers of ECM.^12, 14^ This tissue repair program includes the genes FIZZ1, which drives fibroblast deposition and cross-linking of collagen and is essential for tissue repair in the skin and the lungs;^21, 51, 52, 66^ CCL24, a chemokine for eosinophils that promotes fibrosis in multiple organs;^67–69^ and Fibronectin (FN1), a key ECM protein involved in tissue repair and fibrosis.^5, 70^ We found that this pro-reparative program is also suppressed *in vivo* as tissue repair progresses and tissue stiffness increases. It is known that macrophages are required for successful resolution of tissue repair,^17, 22^ and these findings suggest that macrophage sensing of ECM mechanics could provide the necessary source of negative feedback to maintain ECM homeostasis and prevent unrestrained tissue repair.

Macrophages are able to activate distinct transcriptional programs known as polarization states in order to participate in diverse biological processes. The polarization state involved in tissue repair is traditionally known as the ‘M2’ state and can be induced *in vitro* by the cytokines IL-4 or IL-13. When ‘M2’ macrophages are identified *in vivo*, however, they often express some but not all of the classic marker genes induced by ‘M2’ polarization *in vitro*. This disconnect is poorly understood but may reflect the fact that macrophages respond not only to polarizing signals like IL-4, but also to other information within the tissue environment, as has been suggested by others.^71^ In response to these cues, macrophages may tune their polarization state to meet the specific functional demands of the tissue.^7^ Consistent with this model, we found in this study that a subset of IL-4-induced genes are regulated by key features of the tissue environment—the mechanical properties of the ECM and the abundance of CSF1—while others are insensitive to these inputs. Furthermore, we determined that these environmental inputs are converted to site-specific changes in chromatin availability. This offers an example of separable programs within the ‘M2’ polarization state that are tunable in response to the tissue environment.

This work has uncovered a mechanism of mechanosensing by macrophages that is fundamentally distinct from adhesive cell types, in which ECM mechanosensing has traditionally been studied. Whereas cells such as fibroblasts anchor to the extracellular matrix using integrin receptors and exert pulling forces using non-muscle myosin II to measure the stiffness of the ECM, we found that macrophages in 3D environments do not require integrin-based adhesions or non-muscle myosin II function in order to sense ECM mechanics. This distinction maps onto the differences between the mesenchymal mode of migration used by adhesive cell types like fibroblasts and the amoeboid mode of migration adopted by leukocytes in 3D environments. Thus, we can conceptualize the classical, integrin-dependent mode of mechanosensing as ‘mesenchymal mechanosensing’ and the integrin-independent mode of mechanosensing described here as ‘amoeboid mechanosensing’.

The mechanical properties underlying amoeboid migration remain an active area of investigation. However, one fundamental distinction is that, while mesenchymal migration depends on binding to the ECM and pulling to achieve locomotion, amoeboid migration does not require adhesion and instead depends instead on actin filaments pushing outward at the cell membrane against compressive forces.^39–41, 72–75^ Rather than sensing mechanical tension, therefore, macrophages are likely to sense mechanical compression as they migrate within the tissue. In other words, macrophage mechanosensing may work by pushing, rather than pulling.

It is noteworthy that we made these findings in a 3D tissue environment. Concurrent work has shown that macrophages and T cells can sense ECM stiffness on two-dimensional (2D) surfaces.^42, 46, 76^ In this context, macrophages attach to the underlying ECM and likely use mechanosensing mechanisms that are more similar to adhesive cell types. Indeed, the seminal studies on integrin-independent leukocyte migration within 3D tissues also showed that leukocytes do require integrins in order to migrate on 2D surfaces.^39^ Recent work has further shown that macrophages can utilize both integrin-dependent and integrin-independent modes of migration in 3D environments, depending on ligand availability and the stimulus for migration.^77^ Thus, macrophages may employ different mechanosensing strategies—amoeboid or mesenchymal—depending on the tissue context and whether they are sitting upon or embedded within the extracellular matrix. Indeed, while the majority of macrophages reside in 3D spaces, osteoclasts, alveolar macrophages, and Kupffer cells effectively occupy 2D surfaces and may thus rely on a mesenchymal mechanosensing strategy, similar to other adhesive cell types. Meanwhile, the amoeboid mode of mechanosensing described here may be utilized by other leukocytes, as well as other cell types capable of amoeboid migration,^75^ within interstitial spaces.

Finally, we identified the cytoskeleton as a signal integrator for macrophages. We found that changes in the actin cytoskeleton not only mediate macrophage mechanosensing but are also required for CSF1 to regulate a subset of its genetic targets. This result was surprising. In the case of mechanosensing, it is somewhat intuitive that extracellular structural or mechanical information (in the ECM) would be translated to intracellular structural information (in the cytoskeleton) before being converted to a biochemical signal to regulate gene expression. In the case of growth factor signaling, however, it is not intuitive why an extracellular biochemical signal (CSF1) would act through a cytoskeletal intermediary before being converted back to biochemical information. Yet, this model also appears elsewhere in biology. The best-elucidated example is the regulation, in fibroblasts and other mesenchymal cells, of the transcription factor Serum response factor (SRF). Growth factors present in fetal calf serum induce polymerization of G-actin to F-actin, which causes the transcriptional co-activator MRTF, normally sequestered in the cytoplasm by G-actin monomers, to be released to the nucleus, where it forms a complex with SRF and initiates transcription.^78–81^ Thus, as in CSF1 stimulation of macrophages, a biochemical signal induces cytoskeletal remodeling as an intermediate step to control gene expression.

Importantly, in this case, as in macrophages, these cytoskeletal dynamics are also sensitive to ECM mechanics: Stiff substrates induce increased actin polymerization in fibroblasts, as well, causing MRTF to be released to the nucleus.^65^ Thus, our findings appear to be an example of a broader biological theme, in which the cytoskeleton acts as an integrator of diverse types of information—both mechanical and biochemical—in order to control gene expression programs.

We can speculate as to why the cytoskeleton might have evolved to act as a signal integrator. The cytoskeleton is one of the most ancient features of eukaryotic cells, is essential for the cell’s interaction with its environment, and is critical for a wide range of fundamental cellular activities, including migration, cell division, phagocytosis, protein secretion, and cellular organization within tissues. Cells may have evolved the strategy of measuring cytoskeletal dynamics as a key indicator of both the external state of the tissue and the internal state of the cell, in order to appropriately regulate gene expression programs that need to respond to these cues. Another advantage of using the cytoskeleton as a signaling hub is that it may allow for unique mechanisms of transcriptional regulation that cannot be achieved through canonical signal transduction pathways. For instance, we found here that changes in cytoskeletal dynamics ultimately control chromatin availability. This may take place through mechanical effects of the cytoskeleton on the nuclear membrane^82, 83^ or through regulation of nuclear actin,^84–86^ offering pathways for targeted regulation of mechanosensitive and other cytoskeleton-dependent gene expression programs.

This study also raises fundamental questions for future study, including precisely how extracellular mechanical information is translated into intracellular changes in the cytoskeleton, what specific properties of the actin cytoskeleton are measured by macrophages, and how these changes ultimately control chromatin availability. More broadly, the role of the cytoskeleton in integrating biological information and regulating diverse cellular functions is likely to represent a promising area for future exploration across biological fields.

## Materials and methods

### Cell culture

#### BMDMs

BMDMs were differentiated from mouse bone marrow cells in macrophage growth media (MGM) composed of 30% L929-cell conditioned media and 70% complete RPMI media (cRPMI), which consisted of RPMI-1640 (Corning) supplemented with 10% heat-inactivated fetal bovine serum (FBS) (Gibco), 2 mM L-glutamine, 200 U/mL penicillin/streptomycin, 1 mM sodium pyruvate, and 10 mM HEPES. Bone marrow was prepared by crushing C57BL/6J mouse femurs and tibias to release marrow, followed by ACK (ammonium-chloride-potassium) lysis of red blood cells and passage through a 70-μm cell strainer. Bone marrow was plated at 5 × 10^6^ cells in 20 mL of MGM in a non-tissue culture (non-TC) dish (Day 0). Every three days, cells were supplemented with 10 mL of MGM. After day 6, adherent cells were lifted with cold phosphate-buffered saline (PBS) containing 5 mM EDTA and used for experiments. Bone marrow cells derived from Piezo1^f/f^ and LysM^Cre/+^ Piezo1^f/f^ mice were kindly provided by R. Flavell (Yale University).^45^

#### MEFs

E13.5-E14.5 embryos were collected from a pregnant female by removing the uterus and separating each embryo from its amniotic sac. The head and ‘‘red tissue,’’ including fetal liver, were removed and discarded. The remaining portion of each embryo was minced using razor blades in 0.05% trypsin + EDTA and placed in a 37°C incubator for 30 minutes. After digestion, the tissue was washed and plated in complete DMEM (DMEM + 10% FBS, 2 mM L-glutamine, 1 mM sodium pyruvate, 10 mM HEPES, 200 U/mL penicillin/streptomycin) in 15 cm tissue culture (TC) plates overnight. The following day, cells and undigested tissue debris were lifted from the plates using 0.05% trypsin + EDTA (Gibco) and filtered over a 70-μm cell strainer. These cells were expanded for 1-2 passages and then sorted for CD45-, CD11b-, and F4/80-negativity to exclude contaminating macrophages. The sorted MEFs were split once after sorting to allow for recovery and used for experiments between passage 3 and passage 7.

### Reagents

For stimulation of BMDMs, recombinant murine IL-4 (Peprotech, 214-14) was used at 20 ng/mL, and recombinant murine MCSF (Cell Signaling Technologies, 5228) at 100 ng/mL for 2D culture and 200 ng/mL for 3D culture. Chemical inhibitors used in cell culture included latrunculin A (Cayman Chemical, 10010630 or Sigma, 428021), (-)-blebbistatin (Sigma, B0560), and nocodazole (Sigma, M1404), which were used at 1 μM, 25 μM, and 10 μM, respectively. K-975 (Sellek Chemicals, E1329), RN-1734 (Cayman Chemical, 24205) and HC-067047 (Cayman Chemical, 20927) were all used at 10 μM. GsMTx4 (Abcam, ab141871) was used at 5 μM.

For antibody blockade of cell-surface receptors, cells were pre-incubated in a small volume of media with 160 μg/mL of antibody for 15 minutes at room temperature, followed by dilution to a final concentration of 20 μg/mL for the duration of the experiment. β1 integrin blockade was performed with Purified NA/LE Hamster Anti-Rat CD29 (clone Ha2/5, BD Pharmingen, 555002) and the isotype control antibody Purified NA/LE Hamster IgM, λ1 Isotype Control (clone G235-1, BD Pharmingen, 553957). β2 integrin blockade was performed with Purified NA/LE Rat Anti-Mouse CD18 (clone GAME-46, BD Pharmingen, 555280) and the isotype control antibody Purified NA/LE Rat Ig1, κ Isotype Control (clone R3-34, BD Pharmingen, 553921).

### Three-dimensional collagen gels

#### Preparation of 3D collagen gels with embedded cells for cell culture

Three-dimensional collagen gels with embedded BMDMs or MEFs were synthesized using high-concentration rat tail collagen I (Corning, 354249), using a protocol adapted from Corning’s “alternate gelation procedure” provided with the product. Briefly, Collagen was mixed with 10× PBS and sterile H_2_O to achieve the desired collagen concentration. Subsequently, 1 N NaOH was added to neutralize the acetic acid in the collagen solution until a pH of 7-7.5 was achieved, measured by pH strip. BMDMs or MEFs were added to the collagen solution at 0.5-1 × 10^6^ cells or 0.2 × 10^6^ cells per gel, respectively. Cells were added such that they made up 1/8 of the volume of the gel. Collagen-cell solutions were then plated at 200 ul in 48-well non-TC plates. Until this step, collagen solution was kept on ice as much as possible. After collagen-cell solutions were plated, they were incubated for 45 minutes to 1.5 hours at 37°C and 5% CO_2_ to achieve gelation. A small pipette tip was used to score around the edge of the gels to detach them from the walls of the wells and ensure that they float, in order to avoid cells interacting with the stiff bottom of the plate. Media (400 or 500 ul) was then added, including any stimulation or inhibitors at the appropriate concentrations, and the gels were scored once again before placing them back in the incubator for the duration of the experiment.

#### Harvesting RNA for qPCR from cells in 3D collagen gels

After stimulation for 16-28 hours, gels were transferred with curved, blunt, serrated forceps to a 24-well non-TC plate, rinsing forceps in PBS between handling each gel. RNA-Bee (Tel-Test) 1 ml was then added to each gel and allowed to sit for 5 minutes or until gels began to fragment. RNA-Bee and gel were pipetted up and down until the gel disintegrated and the solution was homogenous. This solution was processed for RNA purification.

#### Harvesting protein for Western blot from cells in 3D collagen gels

Using curved, blunt, serrated forceps, gels were transferred to 200 ul of 2× SDS-PAGE sample buffer with β-mercaptoethanol and protease/phosphatase inhibitor that was pre-heated at 105°C. Samples were further heated at 105°C for 15 minutes, visualized to ensure homogenization, and used for Western blot.

#### Harvesting cells from 3D collagen gels for flow cytometry

Collagenase Type IV (Worthington, LS004189) was added at 18 mg/ml, 100 μl per well, to 500 μl media in which cells were cultured, for a final concentration of 3 mg/ml. Gels were diced with clean scissors directly in the wells. After plates were rotated at 205 rpm for 20 minutes, samples were transferred to conical tubes through a cell strainer, using the top of a plunger to completely mash the partially digested gels. Following 2 washes with PBS, cells were collected with the centrifugation at 1350 rpm (385 x g) at 4°C for 5 minutes.

#### Confocal microscopy of cells in 3D collagen gels

In preparation for confocal microscopy, 200 μl collagen gels were prepared in chambers with coverglass well-bottoms (Lab-Tek Chambered #1.0 Borosilicate Coverglass System, 8-chamber). Gels were not scored so that they would sit on the bottom of the well to be as close as possible to the microscope objective. After gelation, 300 uL of media with recombinant IL-4 and/or CSF1 was added. At the end of the experiment, media was removed with a pipette, gels were washed gently with PBS, and gels were fixed in 300 μl of 4% paraformaldehyde (PFA) for 1 hour at room temperature on a bench-top rotator. Gels were washed three times with 400 μl of PBS and then permeabilized with 0.5% Triton X-100 for 1 hour at room temperature on the rotator. Gels were washed three more times with PBS and incubated with Phalloidin-Texas Red (ThermoFisher Scientific, T7471) at 1:50 in PBS/5%BSA for 1 hour in the dark at room temperature on the rotator. Gels were then counterstained with DAPI in PBS for 5 minutes. Gels were washed three more times with PBS, rotating for 5 minutes in the dark for each wash. Gels were kept in PBS and imaged by Leica SP8 confocal microscopy, scanning for cells that do not contact the bottom of the gel and obtaining Z stacks to capture several whole cells in each image, which were then used for downstream analyses. Cellular morphology was quantified using Imaris 9.8 software (Oxford Instruments) for all cells captured in the acquired confocal images. “Surfaces” were created automatically in the phalloidin channel using the default setting of 0.359 um for “Surfaces Detail” to avoid bias and were then filtered manually to include only genuine cells and only cells whose borders were fully captured. The software was then used to calculate cell sphericity.

### Polyacrylamide-fibronectin (PA) gels

#### Polyacrylamide gel fabrication

Hydrogels were generated by polymerizing various ratios of solutions of acrylamide (Bio-Rad, 3% and 10% for low- and high-stiffness, respectively) and bis-acrylamide (Bio-Rad, 0.3% and 0.225% for low- and high-stiffness, respectively) with ammonium persulfate (Sigma) and tetramethylethylenediamine (Bio-Rad) sandwiched between two glass coverslips. One coverslip was activated with 3-aminopropyltriethoxysilane using 0.1 M NaOH (Macron Fine Chemicals) and 0.05% glutaraldehyde (Polysciences Inc.) and the other coated with dichloromethylsilane (Sigma). Fibronectin was conjugated to the polymerized hydrogels by succinimide chemistry. Briefly, 0.2 mg/mL Sulfa-SANPAH (ThermoFisher Scientific) in 50 mM HEPES (Sigma) were placed under UV (365 nm, 10 mW/cm), washed thoroughly with HEPES, and incubated overnight with 0.2 mg/mL fibronectin (Millipore) at 37°C. After incubation, the hydrogels were washed and kept at 4°C until cell seeding.

#### Cell culture on Polyacrylamide-fibronectin gels

Gels (attached to coverslips) were transferred to 6-well non-TC plates using sterile, curved, pointed forceps. BMDMs (0.85 × 10^6^ cells per gel in 250 μl cRPMI) were gently plated at the center of each gel such that the media formed a meniscus and did not spread beyond the border of the gel. Cells were left to incubate undisturbed for 30 minutes to 1 hour, after which adhesion was confirmed by microscopy. An additional 1.75 ml cRPMI was then added to each gel gently such that the cells were not disrupted. Cell culture plates were then returned to the incubator for 3 hours, after which the stimulation with IL-4 was performed. Cells were then incubated for an additional 16 hours prior to RNA collection.

#### RNA isolation from Polyacrylamide-fibronectin gels

Gels were transferred to new 6-well non-TC plates, taking care to cause minimal mechanical disruption. RNeasy RLT buffer (QIAGEN) with 1:100 β-mercaptoethanol was added to the center of the gel at 350 μl per gel. Several minutes later, RLT buffer on each gel was gently pipetted up and down and collected. RNA was isolated with the RNeasy Micro Kit (QIAGEN) according to manufacturer instructions.

### Scanning Electron Microscopy (SEM)

Collagen gels (400 μl) were formed in 24-well plates and incubated in cRPMI media overnight prior to fixation in 4% PFA for 10 minutes. Hydrogels were then snap-frozen in liquid nitrogen and lyophilized overnight. Freeze-dried samples were sputter coated with palladium and imaged via SEM (Hitachi SU-70). Images were taken at 10 kV for pore quantification. The width of individual pores was analyzed from SEM images using ImageJ, quantifying the pore size as the distance between fibers in three-dimensional space.^87^

### Rheology

A PA or collagen hydrogel was cast between the rheometer base plate and 25 mm diameter parallel plate. Gels were swollen overnight and kept hydrated during testing. The shear modulus of the gel was measured using a strain amplitude sweep of 0.1-10% strain at a constant frequency rate (1 rad/s). The measured shear modulus remained constant over the specified strain range. The elastic modulus was calculated using the shear modulus values, assuming that the gels were incompressible (Poisson ratio = 0.5).

### RNA isolation, cDNA synthesis, and quantitative PCR

After cell collection in RNA-Bee, RNA was isolated according to manufacturer’s instruction, except when isolating RNA from PA gels as indicated. To synthesize cDNA, RNA was annealed to oligo-dT6 primers, and cDNA was reverse-transcribed with MMLV reverse transcriptase (Clontech). Quantitative reverse-transcriptase polymerase chain reaction (qPCR) was performed on a CFX96 or CFX384 Real-Time System (Bio-Rad) using PerfeCTa SYBR Green SuperMix (Quanta Biosciences). Relative expression units were typically calculated as transcript levels of target genes relative to Rpl13a. Primers used for qPCR are listed in Table S1.

### Live cell imaging

Image acquisition was performed with Leica AF6000 Modular System with stage-top incubator INUBTFP-WSKM-F1 (Tokai Hit) maintained at 37°C and 5% CO2. Multiple images were acquired from each sample every 3-5 minutes for 16-20 hours, and videos were assembled from serial images at a single position. Imaris 9.8 was used to track individual cells throughout the video, in order to quantify the max track speed and track displacement length of each cell. “Brownian Motion” was used for the tracking algorithm, in which max distance and max gap size were set as 5 μm and 5, respectively.

### Flow cytometry

All staining steps and washes before cell fixation and permeabilization were performed in FACS buffer (PBS with 2% FBS, 0.01% sodium azide). Cells were incubated with anti-CD16/CD32 (clone 93, eBioscience, 14-0161-86) for Fc receptor blockade and stained with Zombie Yellow (Biolegend) for 10 minutes at room temperature in the dark to distinguish live and dead cells. For lung cells, cell surface markers were stained with the following antibodies: anti-CD11c-FITC (clone N418, eBioscience, 11-0114-85), CD64-Brilliant Violet 421 (clone X54-5/7.1, Biolegend, 139309), Ly6G-PerCP-Cy5.5 (clone 1A8, BD Biosciences, 560602), F4/80-PE/Cyanine7 (clone BM8, Biolegend, 123113), CD45.2-BUV395 (clone 104, BD Biosciences, 564616), and CD11b-BUV737 (clone M1/70, BD Biosciences, 612800). After washing, cells were fixed and permeabilized with BD Cytofix/Cytoperm for 15-20 minutes. Cells were then washed with BD Perm/Wash, and intracellular staining was performed in BD Perm/Wash for 45 minutes at room temperature or at 4°C overnight. Antibodies included anti-FIZZ1-PE (clone DS8RELM, eBioscience, 12-5441-80) and ARG1-APC (clone A1exF5, eBioscience,17-3697-82). Rat IgG1κ-PE (clone R3-34, BD Biosciences, 554685) and Rat IgG2aκ-APC (clone eBR2a, eBioscience, 17-4321-81) were also used as isotype control for gating FIZZ1- and ARG1-positive cells, respectively. After washing with BD Perm/Wash, cells were run on a BD LSR II Green, followed by analysis with FlowJo software (FlowJo LLC).

### Western blot

Samples were run on Bio-Rad Mini-PROTEAN TGX Stain-Free Gels, 4-15%, with a 10- or 12-well comb in Tris/Glycine/SDS buffer. Protein was transferred onto activated PVDF membranes using the Bio-Rad Trans-Blot Turbo system according to manufacturer instructions. Membranes were blocked with TBST (20 mM Tris, 150 mM NaCl, 0.05% Tween 20)/5% BSA for 1 hour and then incubated with primary antibodies in TBST/5% BSA with sodium azide at 4°C overnight. Primary antibodies included rabbit anti-phospho-STAT6 (Cell Signaling Technologies, 56554), rabbit anti-RELMα (Peprotech, 500-P214), and mouse anti-GAPDH (Santa Cruz, sc-32233). Membranes were washed three times with TBST, then incubated with horseradish peroxidase-conjugated secondary antibodies (anti-rabbit or anti-mouse) for one hour at room temperature. Membranes were washed three more times and then developed using Pierce ECL Western Blotting Substrate (ThermoFisher Scientific) or SuperSignal West Pico Chemiluminescent Substrate (ThermoFisher Scientific). Protein was visualized using the Bio-Rad Image Lab system.

### Talin1 knockdown

Gene knockdown was performed using P2 Primary Cell 4D-Nucleofector^TM^ X Kit (Lonza) according to manufacturer’s instruction with modification. Briefly, 2 × 10^6^ BMDMs in a cuvette was transfected with 500 pmol of MISSION® siRNA Universal Negative Control #1 or TLN1 siRNA (ID: SASI_Mm01_00114842) (Sigma) with the protocol “Mouse, macrophage” programmed in 4D-Nucleofector^TM^ (Lonza). Cells were cultured in the media composed of 50% L929-cell conditioned media and 50% cRPMI. Two days later, cells were harvested and used for 3D culture.

### BMDM culture on TC plate

BMDMs were seeded at 0.33 × 10^6^ cells per well of a 12-well dish. After stimulation for 20-24 hours, cells were photographed using a Zeiss Axio Vert.A1 microscope for cell morphology analysis, lysed in RNA-Bee for mRNA expression analysis, or detached with PBS + 5 mM EDTA as described above for flow cytometry analysis.

### RNA sequencing (RNA-seq) and analysis

The aqueous phase after chloroform extraction from RNA-Bee solution was mixed 1:1 with 70% ethanol and further processed with QIAGEN RNeasy Mini columns, with on-column DNase treatment, according to manufacturer instructions. Sequencing libraries were constructed following Illumina Tru-seq stranded mRNA protocol. Paired-end sequencing was performed with Next-seq 500 with paired end reads of 38 base pairs. Illumina fastq files were downloaded from Illumina Basespace and were aligned with Kallisto v0.46.1 using the “-b 100 and -t 20” options to obtain transcript abundances in transcripts per million (TPM) and estimated counts.^88^ The kallisto index used during transcript quantification was built from the Mus musculus transcriptome GRCm38 downloaded as a fasta file from Ensembl (ftp://ftp.ensembl.org/pub/release-90/fasta/ mus_musculus/cdna/s). Transcripts were annotated using the Bioconductor package biomaRt v2.40.5.^89^ Significant differences in gene expression between conditions were calculated, with correction for multiple comparisons, using Sleuth in R v3.5.1.^88^

### Assay for transposase-accessible chromatin with high-throughput sequencing (ATAC-seq)

ATAC-seq was performed according to the protocol detailed by Buenrostro and colleagues.^90^ UCSC Genome Browser (https://genome.ucsc.edu/) was used for data visualization. HOMER was used to analyze known transcription factor motif enrichment (http://homer.ucsd.edu/homer/). Paired-end sequencing was performed with Next-seq 500 with paired end reads of 38 base pairs. Illumina fastq files were downloaded from Illumina Basespace, and adapters were removed with Cutadapt (https://journal.embnet.org/index.php/embnetjournal/article/view/200) using the option “-a CTGTCTCTTATA -A CTGTCTCTTATA -m 23”. Reads were then mapped to the mm9 mouse genome with Bowtie2 using the option “-k 10 -X 1000 –end-to-end –very-sensitive”.^91^ Sam files were converted to Bam files, and read duplicates were removed using Picard (https://broadinstitute.github.io/picard/). Blacklist regions (obtained from https://github.com/MayurDivate/GUAVA) and mitochondrial genomic reads were removed using BEDTools (https://academic.oup.com/bioinformatics/article/26/6/841/244688) and Picard, respectively. Bam files were converted to Bed files also using BEDTools, and macs2 (https://genomebiology.biomedcentral.com/articles/10.1186/gb-2008-9-9-r137) was used to call peaks using the converted bed file. Macs2 call-peak options used were “-f BED -g mm -q 0.05 -- nomodel --shift -75 --extsize 150 --keep-dup all”. To count the number of reads in each peak, featureCounts (https://academic.oup.com/bioinformatics/article/30/7/923/232889) was used. Read counts were normalized using the TMM method in EdgeR (https://bioconductor.org/packages/release/bioc/html/edgeR.html) and differences of 1.5-fold or greater were analyzed as discussed in the text.

### Animals

All use of animals was performed in accordance with institutional regulations after protocol review and approval by Yale University’s Institutional Animal Care and Use Committee.

### Bleomycin-induced lung injury

C57BL/6 mice of 8-10 weeks of age were injected oropharyngeally with 50 ul of PBS or bleomycin 1 U/ml (Teva) after being anesthetized with 1,2-propanediol and isoflurane at a ratio of 7:3. On days 3, 6, 9 or 12, mice were sacrificed and transcardially perfused with 60 ug/ml heparin in PBS. For histology analysis, lung was fixed and inflated by intratracheal injection with 10% PFA. Whole lung was collected and further fixed in 10% PFA for 6 hours at 4°C. The left superior lobe was subjected to HE staining and Masson’s trichrome staining, which were performed by Yale Pathology Tissue Services. The right superior lobe was further treated with 30% sucrose for 24 hours and embedded in O.C.T compound (VWR). Frozen sections were prepared and stained as described below. For mRNA expression analysis, all lung lobes were homogenized in 1 ml RNA-Bee, the supernatant was collected after centrifugation at 12,700 rpm for 3 minutes, and RNA isolation and qPCR analysis was performed. For flow cytometry analysis, whole lung was minced with a sharp razor blade and digested with collagenase type IV 2 mg/ml (Worthington, LS004189) and deoxyribonuclease II 4 μg/ml (Worthington, LS002425) at 37°C with 180 r/min of rotation for 30 minutes. Cells were passed through a 70-μm cell strainer and remaining tissue pieces were mashed with a plunger. Following red blood cell lysis by ACK Lysing Buffer (Lonza), single cell suspension was collected and stained as described above. Isotype control antibodies for gating Fizz1- and Arg1-positive cells were used at each time point for each sample.

### Immunostaining of lung tissue

Frozen lung tissues were cut at 60 μm thickness with a cryostat. After washing with PBS, sections were incubated with 0.3% Triton-X in PBS (wash buffer) for 30 minutes for permeabilization and then with 5% Normal goat serum (Jackson ImmunoResearch) in wash buffer (blocking buffer). Endogenous biotin was blocked with Avidin/Biotin Blocking Kit (Vector Laboratories) according to manufacturer’s instruction. Following the wash with wash buffer, tissues were reacted with rabbit anti-collagen I antibody (ThermoFisher Scientific, PA5-95137) at 1:200 and biotinylated anti-F4/80 antibody (clone BM8, eBioscience, 13-4801-85) at 1:200 in blocking buffer overnight at 4°C. Sections were washed with wash buffer, incubated with goat anti-rabbit antibody conjugated with Alexa Fluor Plus 488 (ThermoFisher Scientific, A32731) at 1:1000 and streptavidin conjugated with Alexa Fluor 647 (ThermoFisher Scientific, S32357) at 1:2500, and washed again. Subsequently, sections were mounted with ProLong™ Diamond Antifade Mountant with DAPI (ThermoFisher Scientific) and observed under Leica SP8. Z-stack images were taken at 1 μm steps from the bottom of the section to the top. Cell sphericity was calculated in the channel for F4/80 using Imaris 9.8 as described above except for filtering “Number of voxels Img = 1” ≥ 5000 to avoid capturing cell debris. DAPI-positive cells were further selected to only include genuine cells for cell sphericity calculation.

### Statistical analyses

Unpaired t tests and an analysis of variance (ANOVA) test with multiple comparisons were used where appropriate to determine statistical significance between groups (p < 0.05 was considered significantly different) using Prism 9.2 (GraphPad Software, Inc). All data are representative of 2-3 independent experiments and shown as mean ± SD, unless specified. *p < 0.05, ** p < 0.01, *** p < 0.001, **** p < 0.0001. ns; not significant.

## Supporting information

Movie S1. Low collagen

Movie S1. High collagen

Movie S2. Low collagen

Movie S2. Low collagen + CSF1

## Acknowledgments

Work in the R.M. lab is supported by the Howard Hughes Medical Institute (HHMI), the Blavatnik Family Foundation, the Scleroderma Research Foundation, and a grant from the NIH (1R01 AI144152-01). M.L.M. was also supported by the NIH MSTP Training Grant T32GM136651 and NHLBI F31 predoctoral fellowship HL139116-01A1. Y.K. was supported by The Uehara Memorial Foundation and Japan Society for the Promotion of Science (JSPS). Electron microscopic images were taken by the Center for Cellular and Molecular Imaging (CCMI) at Yale University. Preparation of lung sections and staining were performed by Yale Pathology Tissue Services. Image analysis was performed at the *In Vivo* Imaging Facility of Yale University. Cells and reagents and were generously provided by Richard Flavell (Yale University), Gwendolyn Randolph (Washington University in St. Louis), David Critchley (University of Leicester), Megan King (Yale University), and Stephen Smale (University of California Los Angeles). We thank Shuang Yu in R.M. lab for assistance processing lung specimens and Michael Sixt (Institute of Science and Technology Austria) for his thoughtful advice. We would like to thank all the members of the Medzhitov Lab and many other members of the Yale Immunobiology Department for invaluable input and support throughout the course of this work.

