## Supplementary Materials for "Macrophages sense ECM mechanics and growth factor availability through cytoskeletal remodeling to regulate their tissue repair program"

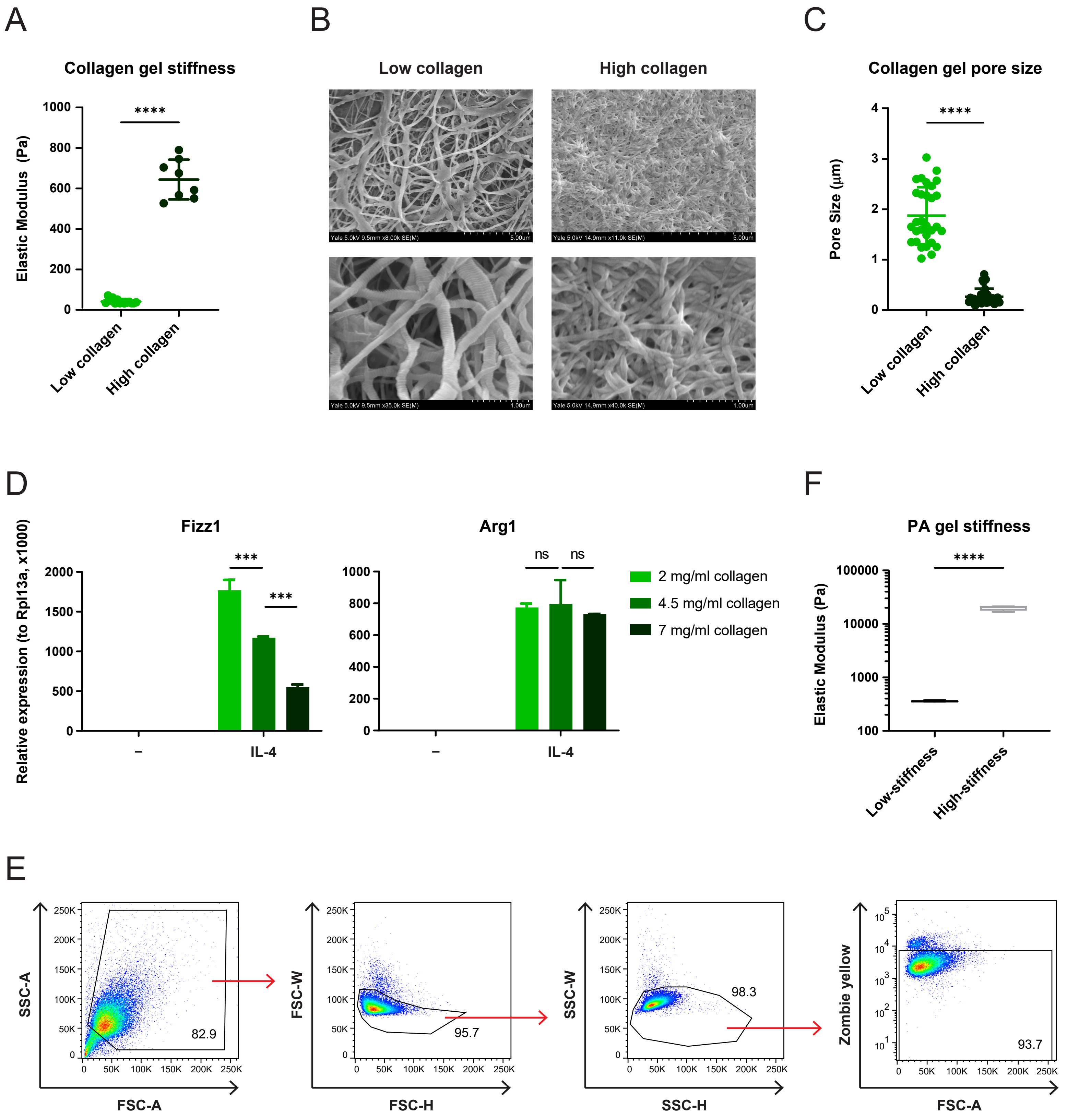


**Fig. S1. Mechanical properties of collagen gels and PA hydrogels**

**(A)** Stiffness of low- and high-collagen gels, measured by rheology. **(B)** Scanning electron microscopy (SEM) images of low- and high-collagen gels at lower (above) and higher (below) magnification. **(C)** Pore sizes of low- and high-collagen gels quantified in SEM images. **(D)** Relative gene expression in BMDMs cultured in indicated concentration of collagen gels with or without IL-4. **(E)** Gating strategy for flow cytometry analysis of FIZZ1 and ARG1 protein expression in BMDM. **(F)** Stiffness of low- and high-stiffness PA gels, measured by rheology. Data are represented as mean ± SD. Student's t test (A, B, and F) and two-way ANOVA (D) are used for statistical analysis. *** $p < 0.001$, **** $p < 0.0001$. ns; not significant.

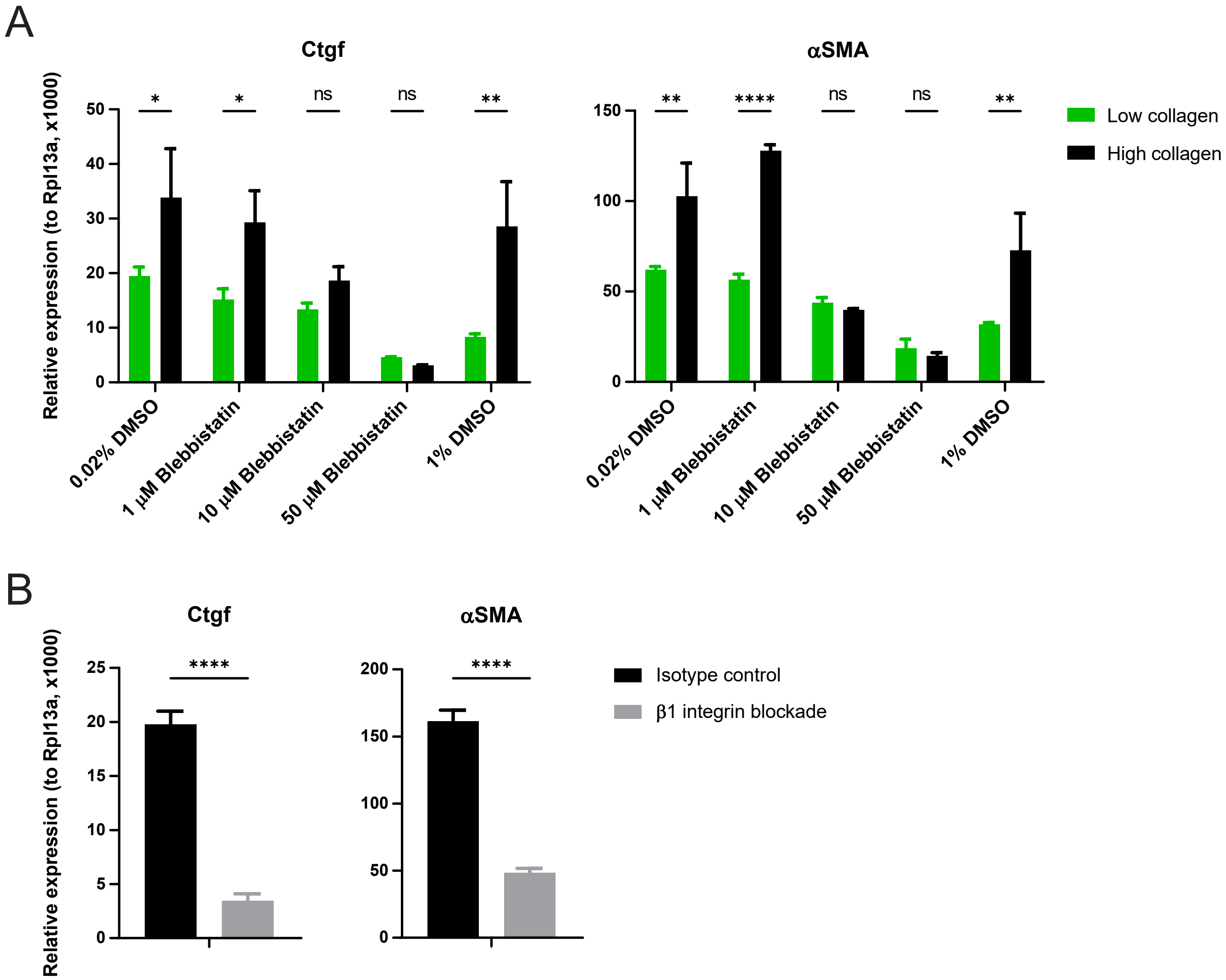


**Fig. S2. Fibroblast mechanosensing is integrin- and non-muscle myosin II-dependent.**

**(A)** Relative gene expression in MEFs cultured in low- or high-collagen gels with the indicated doses of blebbistatin or DMSO (equivalent to the amount of DMSO in the lowest and highest concentrations of blebbistatin). **(B)** Relative gene expression in MEFs cultured in 3D low-collagen gels with β1 integrin-blocking antibody or isotype control antibody. Data are represented as mean ± SD. Two-way ANOVA (A) and Student's t test (B) were used for statistical analysis. *p < 0.05, ** p < 0.01, **** p < 0.0001. ns; not significant.

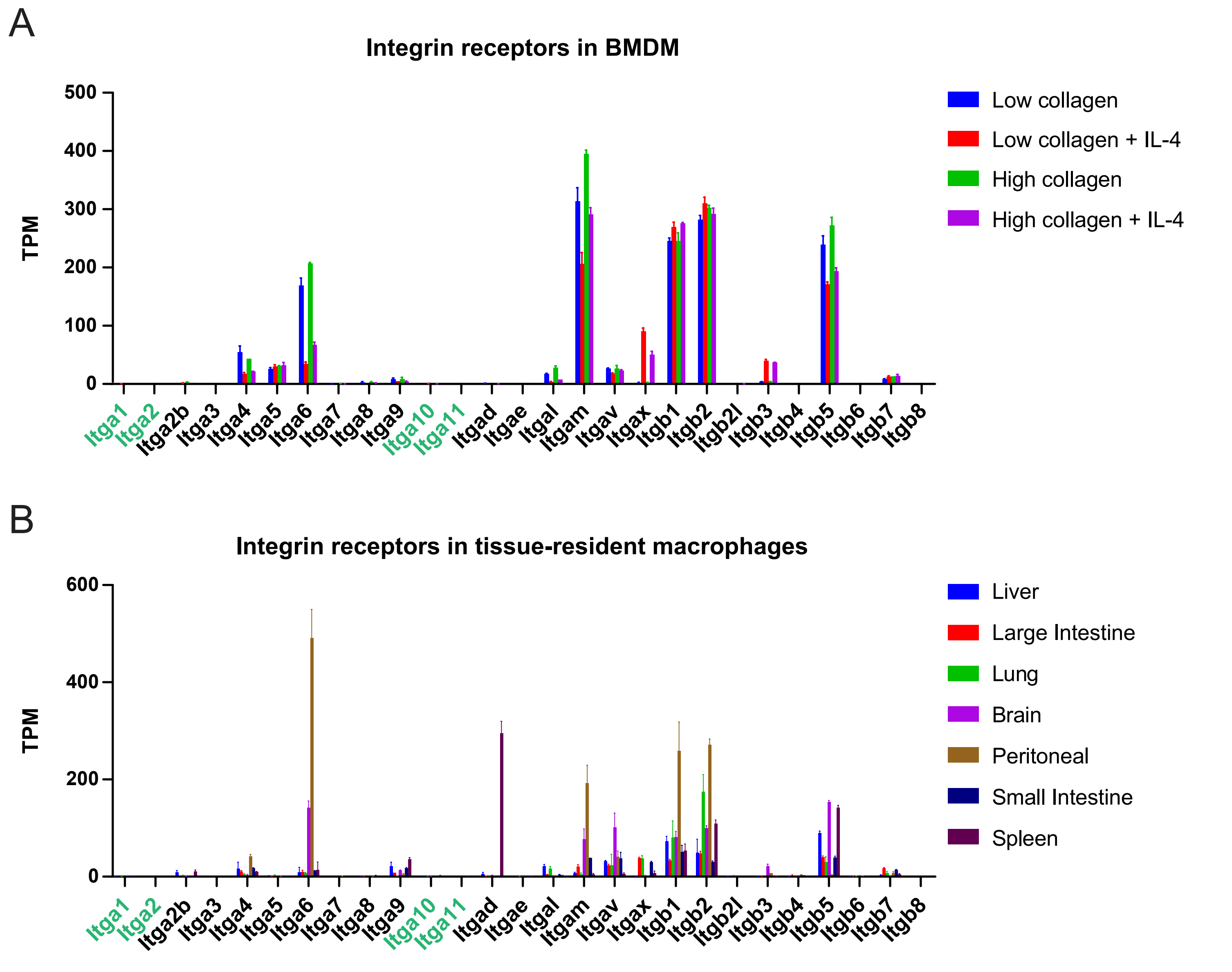


**Fig. S3. Macrophages show minimal expression of collagen-binding integrins.**

**(A)** Normalized gene expression levels (transcripts per million, TPM) of all integrin alpha and beta chains in BMDMs, from RNAseq analysis of BMDMs cultured in 3D low- or high-collagen gels, with or without IL-4 stimulation. **(B)** Normalized gene expression levels (TPM) of all integrin alpha and beta chains in various tissue-resident macrophage populations, based on reanalysis of RNAseq data from Lavin et al.[56] Alpha chains of collagen-binding integrins are highlighted in green (Itga1, Itga2, Itga10, Itga11). Data are represented as mean ± SD.

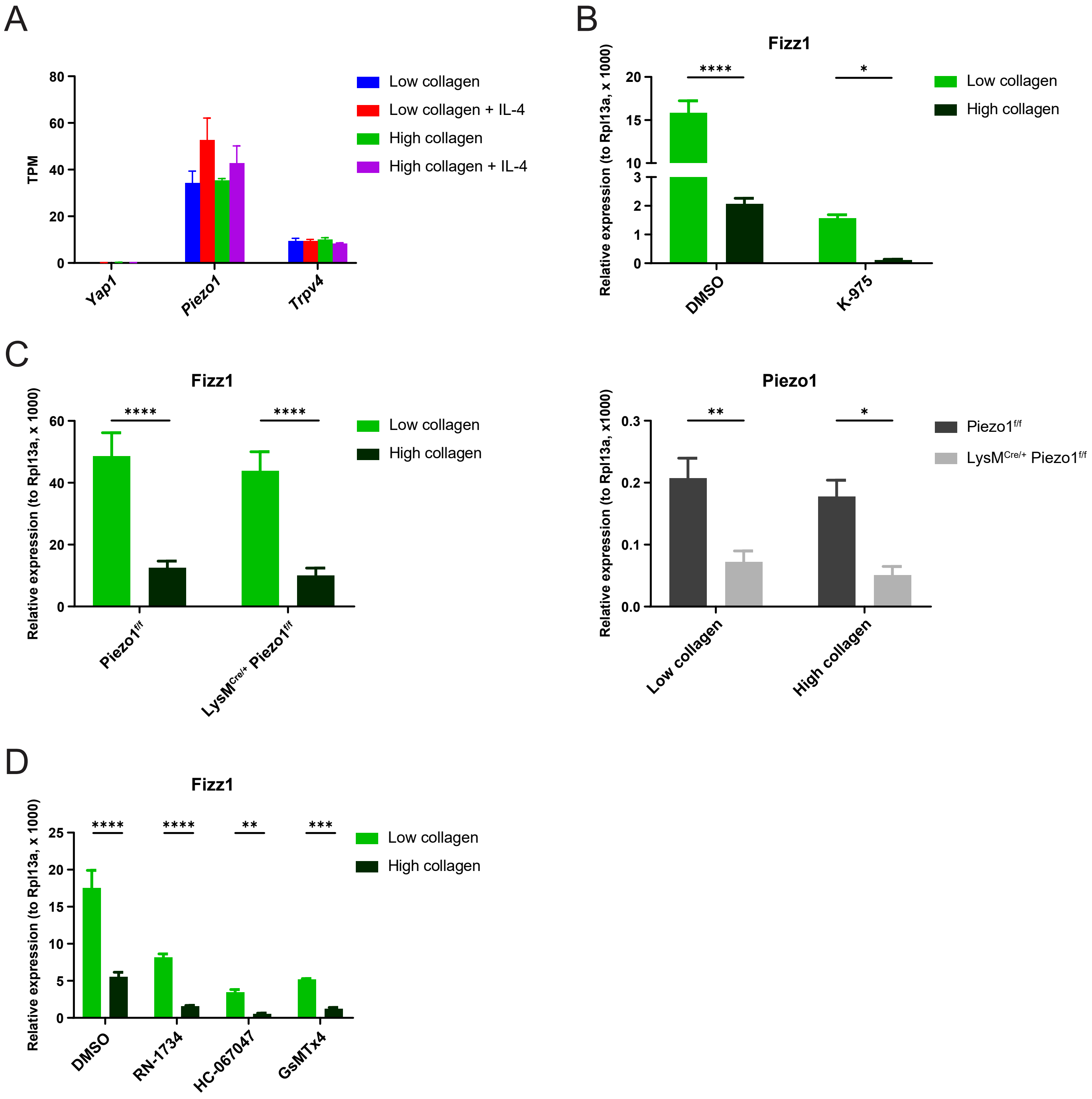


**Fig. S4. YAP signaling and mechanosensitive ion channels are not required for macrophage mechanosensing.**

**(A)** TPM of *Yap1*, *Piezo1* and *Trpv4* from RNAseq analysis of BMDMs cultured in 3D low- or high- collagen gels with or without IL-4 stimulation. **(B)** Relative gene expression in BMDMs treated with control or K-975 (TEAD inhibitor) and cultured in low- or high-collagen gels in the presence of IL-4. **(C)** Relative gene expression in BMDMs derived from Piezo1$^{f/f}$ or LysM$^{Cre/+}$ Piezo1$^{f/f}$ mice, cultured in low- or high-collagen gels in the presence of IL-4. **(D)** Relative gene expression in BMDMs treated with control, RN-1734 or HC-067047 (TRPV4 inhibitors), or GsMTx4 (mechanosensitive ion channel inhibitor) and cultured in low- or high-collagen gels in the presence of IL-4. Data are represented as mean ± SD. Two-way ANOVA was used for statistical analysis. *$p < 0.05$, ** $p < 0.01$, *** $p < 0.001$, **** $p < 0.0001$.

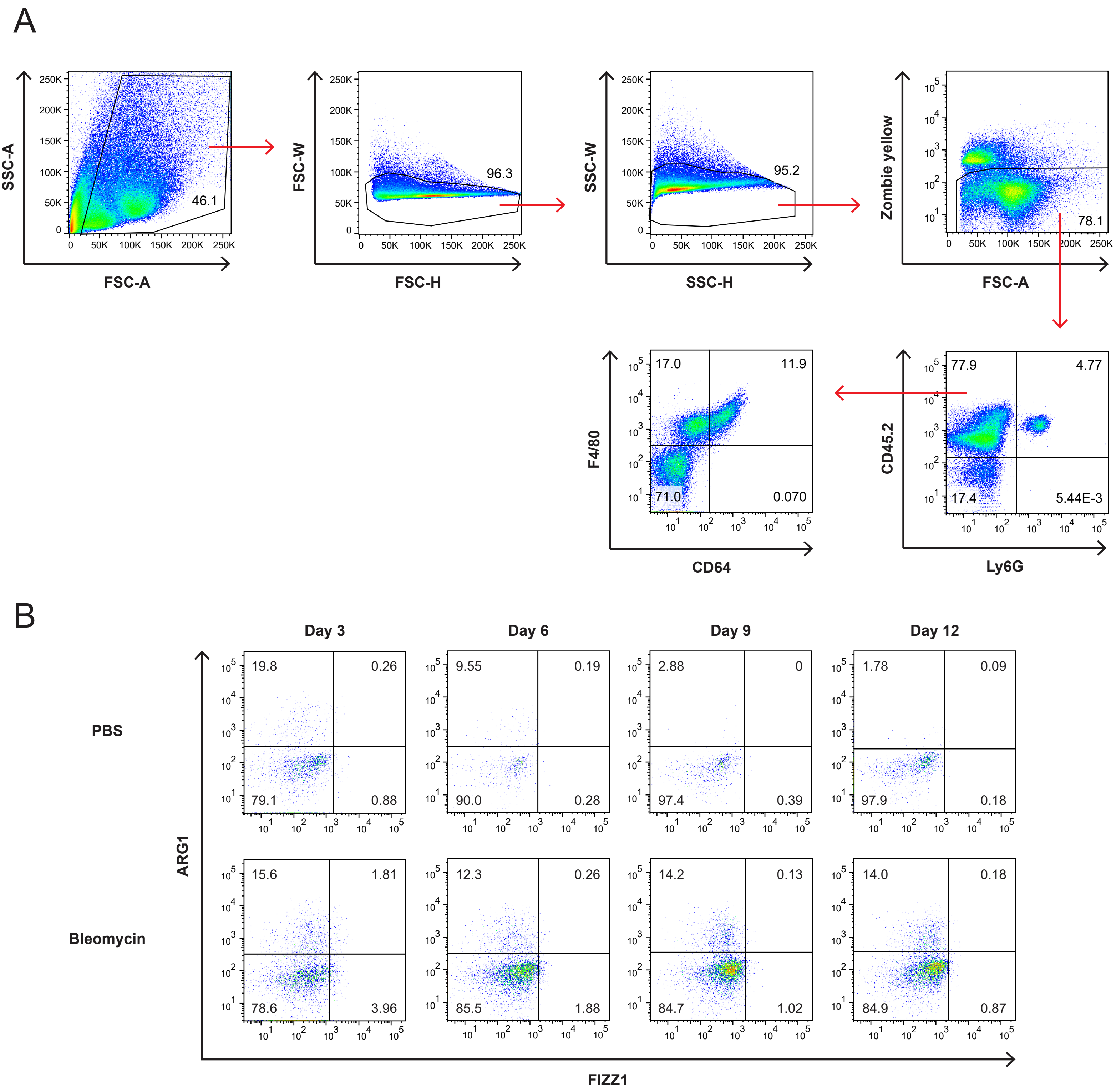


**Fig. S5. Mechanosensitive gene expression program is induced during tissue repair *in vivo*.** **(A)** Gating strategy for flow cytometry analysis of FIZZ1 and ARG1 protein expression in mouse lung macrophages. **(B)** FIZZ1 and ARG1 protein expression in lung macrophages in PBS- or bleomycin-treated mice, analyzed by flow cytometry. Representative plots following the gating strategy in (A) are shown.

Movie S1. Macrophages in 3D low- and high-collagen gels

Movie S2. Macrophages in 3D low-collagen gels with or without CSF1

**Table S1.** List of primers used for qPCR analysis.

| Gene name | | Sequence (5' to 3') |
|---|---|---|
| ***Arg1*** | Forward | CTGGTGTGGTGGCAGAGG |
| | Reverse | TGGCCAGAGATGCTTCCAAC |
| ***Mrc1*** | Forward | AAAGGGACGTTTCGGTGGAC |
| | Reverse | CACTCCGGTTTTCATGGCAAC |
| ***Fizz1*** | Forward | GATGAAGACTACAACTTGTTCC |
| | Reverse | AGGGATAGTTAGCTGGATTG |
| ***Rnase2a*** | Forward | TCCACGGGAGCCACAAAG |
| | Reverse | GAGGCAAGCATTAGGACATGTC |
| ***Ear2*** | Forward | TCCACGGGAGCCACAAAG |
| | Reverse | GAGGCAAGCATTAGGACAAGTC |
| ***Ym1*** | Forward | CCCTACAATTAGTACTGGCCCAC |
| | Reverse | CCTCAGTGGCTCCTTCATTCAG |
| ***Fn1*** | Forward | CAACCTCTGCAGACCTACCC |
| | Reverse | ACTGGATGGGGTGGGAATTG |
| ***Ccl24*** | Forward | AGCATCTGTCCCAAGGCAG |
| | Reverse | TGTATGTGCCTCTGAACCCAC |
| ***Ctgf*** | Forward | AGGGCCTCTTCTGCGATTTC |
| | Reverse | GACCCACCGAAGACACAGG |
| ***Acta2*** | Forward | ATCACCATTGGAAACGAACGC |
| | Reverse | TAGGTGGTTTCGTGGATGCC |
| ***Col1a1*** | Forward | ACGAGATGGCATCCCTGGA |
| | Reverse | GCCATAGGACATCTGGGAAGC |
| ***Talin1*** | Forward | TCTACCATGGTGTACGACGC |
| | Reverse | GACAGAAAGAGCCCAAAGTCG |
| ***Rpl13a*** | Forward | GAAGGAAAAGGCCAAGATGCAC |
| | Reverse | TGAGGACCTCTGTGAACTTGC |
